# Is cultural context the crucial touch? Neurophysiological and self-reported responses to affective touch in women in South Africa and the United Kingdom

**DOI:** 10.1101/2025.01.21.634031

**Authors:** Danielle Hewitt, Sahba Besharati, Victoria Williams, Michelle Leal, Francis McGlone, Andrej Stancak, Jessica Henderson, Charlotte Krahé

## Abstract

Affective touch, involving touch-sensitive C-tactile (CT-) afferent nerve fibres, is integral to human development and wellbeing. Despite presumed cultural differences, affective touch research typically includes ‘Western’, minority-world contexts, with findings extrapolated cross-culturally. We report the first cross-cultural study to experimentally investigate subjective and neurophysiological correlates of affective touch in women in South Africa (SA) and the United Kingdom (UK) using (1) subjective touch ratings, and (2) cortical oscillations for slow CT-optimal (vs. faster non-CT-optimal) touch on two body regions (arm, palm). Cultural context modulated affective touch experiences: SA (vs. UK) participants rated touch as more positive and less intense, with enhanced differentiation in sensorimotor beta band oscillations, especially during palm touch. UK participants differentiated between stroking speeds, with opposite directions of effects at arm and palm for frontal theta oscillations. Results highlight the importance of cultural context in subjective experience and neural processing of affective touch.

---

Our sense of touch is essential for physical and social interaction. Discriminative touch functions to identify object properties and guide motor behaviour ^1^, whereas affective touch – typically gentle, stroking (i.e., dynamic) touch – fulfils affiliative ^2^ and communicative ^3–5^ social functions. Affective touch is typically associated with pleasant feelings ^7^ (although see e.g., ^6^) and approach tendencies ^7^. Research has charted the importance of affective touch, and more broadly prosocial/affectionate touch, for human social development ^8^, emotion regulation ^9^, and psychological and physical wellbeing ^10,11^. However, the empirical studies informing our knowledge of the perception, functions, and effects of affective touch have overwhelmingly been carried out in ‘Western’ contexts (sometimes termed minority-world settings, differentiating these from majority-world contexts, where most of the world’s population resides ^12^), limiting our understanding of cross-cultural variations in touch perception and processing.

The few studies investigating the role of culture in affective touch ^13–16^ have largely focused on comparing self-reported outcomes, with no studies on the neurophysiological correlates of affective touch in different cultural contexts. Given presumed cultural differences in touch norms ^15^ and early touch experiences (e.g., differences in baby wearing and co-sleeping across cultures ^17^), together with the influence of such ‘top-down’ factors on touch perception and evaluation (e.g., ^4^; see ^18^ for a review), it is important to examine whether affective touch is perceived and processed distinctly across cultural contexts. Accordingly, this pre-registered (https://doi.org/10.17605/OSF.IO/FCQNK) experimental study conducted in the United Kingdom (UK) and South Africa (SA) explored how cultural context shapes touch evaluations and neural oscillations (captured using electroencephalography; EEG) during affective touch.

Affective touch perception is mediated by the integration of ‘bottom-up’ peripheral afferent pathways and top-down psychological and contextual factors. Tactile stimulation activates cutaneous low-threshold mechanoreceptors via myelinated Aβ afferent fibres, resulting in rapid central processing through the somatosensory system (reviewed in ^19^). Additionally, gentle stroking of hairy skin at velocities of 1–10cms ^−1^, optimally at 3cms ^−1^, preferentially activates unmyelinated C-tactile (CT) afferents ^20^, with such activation positively correlated with perceived pleasantness ^20,21^. Experimentally, this type of touch is commonly contrasted with faster, non-CT-optimal touch to the hairy skin ^20^ or touch to a non-hairy (glabrous) body region such as the palm (where CT fibres are not – or only sparsely – present) to isolate the contribution of CT-fibre activation and bottom-up from top-down effects on touch perception.

Neuroimaging studies have highlighted a network of brain regions involved in touch processing, including primary and secondary somatosensory cortices, insular cortex, posterior parietal cortex, and orbitofrontal cortex ^22^. Affective, CT-optimal touch (compared to touch at faster velocities or to non-CT-innervated skin) has been associated with stronger activation of the dorsal posterior insula ^23^, a key region for interoception ^24–26^. However, temporal dynamics during touch are better captured with electroencephalography (EEG), which detects real-time changes in the amplitude of cortical oscillations (event-related synchronisation, ERS, and desynchronisation, ERD) during sensory processing. ERD and ERS in alpha (8–13 Hz) and beta bands (16–24 Hz) have been linked with active inhibition or cortical activation in the sensorimotor system, respectively ^27,28^. Tactile brushing stimulation of glabrous and hairy skin elicits ERD in alpha and beta bands over bilateral SI/MI cortices, suggesting their involvement in bottom-up sensory processing and motor preparation ^29^. In contrast, midfrontal theta (4–7 Hz) oscillations are implicated in cognitive control ^30^ and emotion regulation ^31^. Literature on the neural oscillations during affective touch is limited, with only one study showing that CT-optimal touch (to hairy skin) attenuated widespread theta band oscillations and parietal beta oscillations compared to non-CT-optimal touch ^32^, pointing towards a more specialised involvement of these oscillations in emotional and attentional responses during affective touch. However, to our knowledge, oscillatory changes between the arm and palm during tactile stimulation have not been systematically compared.

Top-down psychological factors and individual differences influence the perception and meaning ^18^, as well as the neurophysiological correlates, of affectionate touch ^33,34^. Touch experiences (e.g., the amount of exposure to touch ^35^) and touch attitudes (e.g., as part of attachment styles ^36^) modulate perceived pleasantness of affective touch, while holding a partner’s hand (static touch involving glabrous skin) after a negative affect induction attenuated theta and beta activity, but (for theta) only in people with a secure attachment style ^33^. Moreover, sex and gender effects are apparent, with females rating affective touch more positively ^37^ and women ascribing different meaning to touch ^4^ compared to males/men. At the level of cultural context, individuals from collectivist cultures report higher acceptability of affectionate touch ^13,15^. However, potential differences at the neural oscillatory level remain to be investigated.

Accordingly, in the current study, we experimentally tested evaluative and cortical oscillatory responses to affective touch using a cross-cultural comparison of women living in SA vs. the UK. In a fully within-subjects design, we varied touch velocity (affective i.e., slow, CT-optimal, vs. faster, non-CT-optimal) and body region (arm vs. palm) to tease apart the influence of bottom-up vs. top-down effects. We hypothesised that across cultural contexts, slower-velocity, affective touch would be evaluated more positively than faster-velocity touch ^20^, and would be associated with decreased theta band activity (specifically, increased ERD) in response to CT-optimal touch (cf. ^32^). We also examined effects of affective touch on alpha and beta band activity to capture sensorimotor processes. Given greater innervation of Aβ fibres and relevance of the hand for reach-to-grasp movements, we explored whether increased alpha and beta ERD would be evident for the palm vs. arm. We had no specific hypotheses for body region for theta, given attenuated theta in response to touch at arm and palm regions ^32,33^.

Information on touch norms is sparse, but SA shares some factors associated with more positive touch norms and greater touch frequency (greater collectivist tendencies and warmer climate^13^) compared to the UK. We thus hypothesised that SA participants would evaluate affective touch more positively and, given potentially greater touch exposure in the SA context (see ^35^, for effects of touch exposure on affective touch perception), SA participants would show enhanced differentiation (to slow vs. faster touch) in neural oscillations compared to UK participants. We controlled for individual differences in touch experiences and attitudes, and attachment styles.

Taken together, we report the first experimental study on the influence of cultural context on neurophysiological correlates of affective touch, controlling for individual differences in touch attitudes and experiences. Given the increasing use of touch in therapeutic contexts ^38^, indications of cross-cultural differences in health benefits as a function of affective touch ^39^, and the cultural diversity within and between countries in an increasingly globalised world, it is crucial to understand how touch is experienced at both self-report and neural levels, contingent on the culture-specific meanings and functions of touch.

## Results

### Descriptive statistics

Self-reported ratings of the dynamic, stroking touch (how much the touch was liked, wanted, and how intense, comfortable, and pleasant it was), are presented by country in Table 1 (see Supplementary Table 1 for ratings across countries). There were no differences between SA and UK for any of the individual difference variables (attachment styles and experiences and attitudes to touch; see *Methods*) related to the perception and evaluation of touch, except that UK participants had significantly more positive attitudes to unfamiliar touch than did SA participants (see *Participants*). All statistical analyses reported below controlled for individual differences, as pre-registered. In our pre-registration, we stated that we would primarily examine effects across countries, and investigate effects of country in an exploratory step. We carried out the analyses largely as pre-registered (explicitly noting deviations below). As our data showed a strong modulatory effect of cultural context, which qualified the effects across countries, we here present analyses including effects of cultural context on outcomes and report findings from across-country models in Supplementary Materials.

**Table 1.**
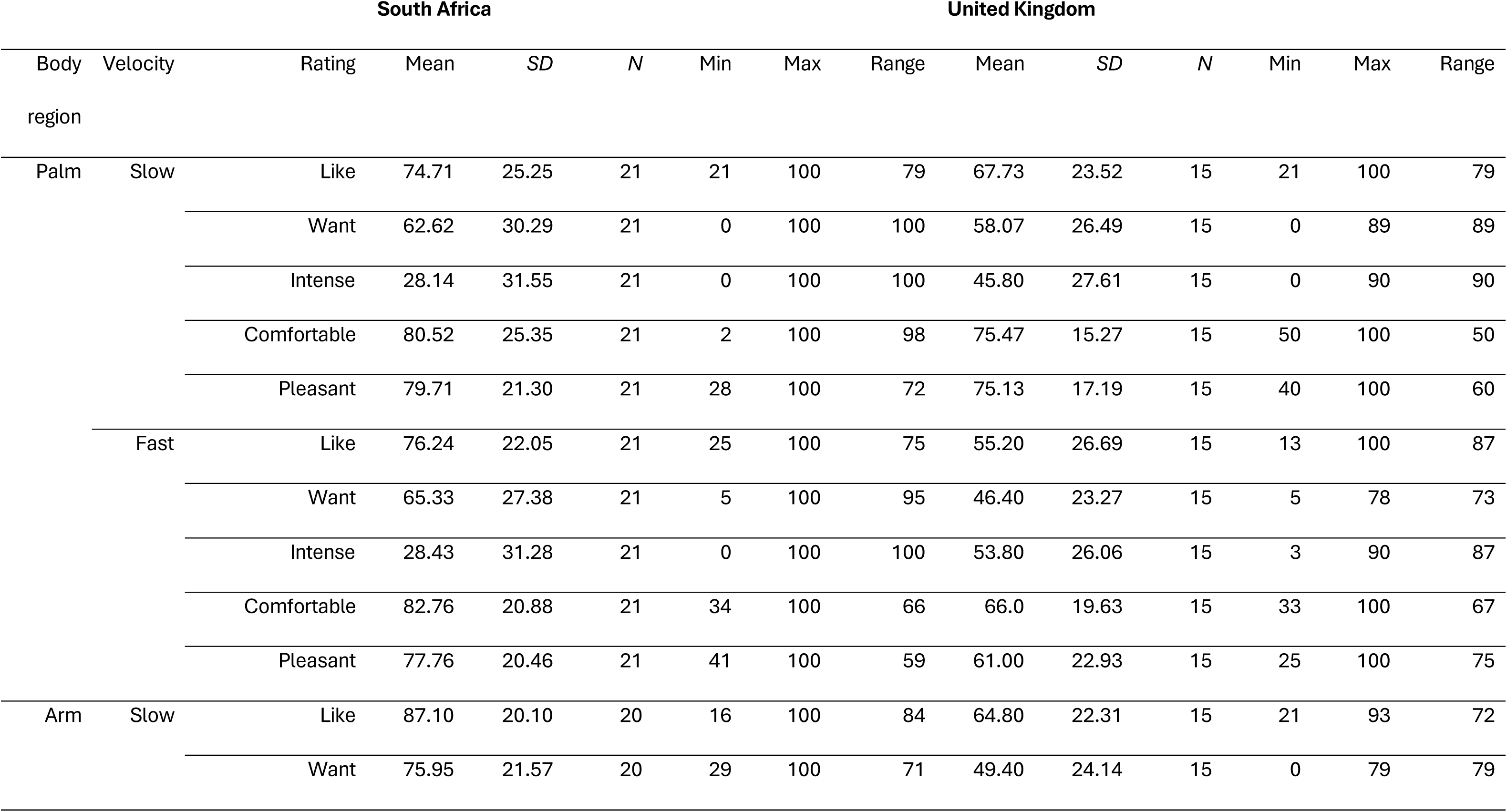
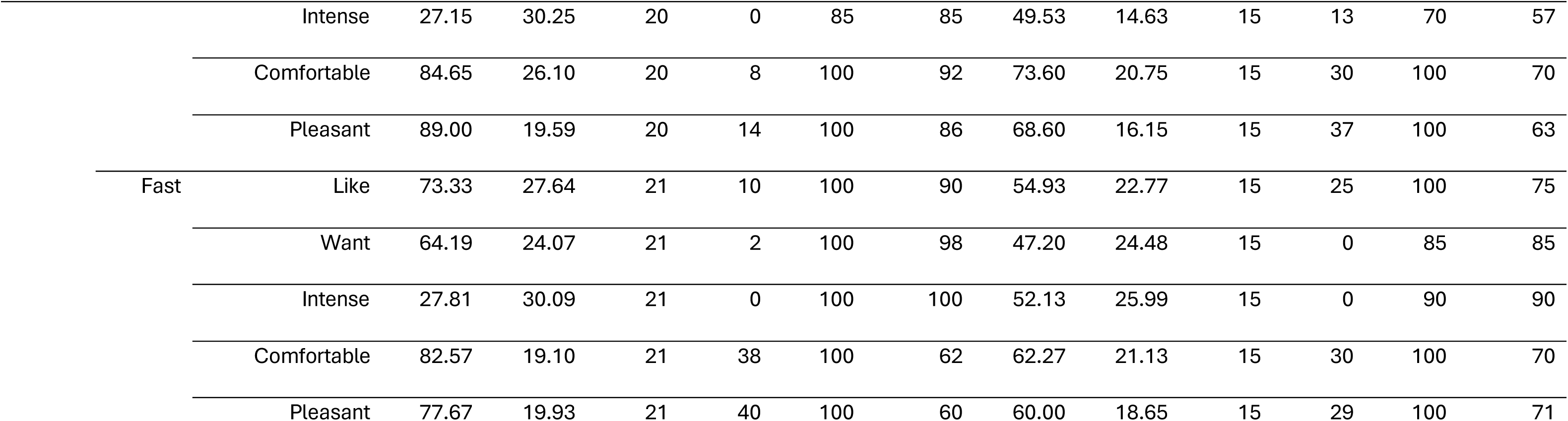
Descriptive statistics for touch ratings by country.

### Touch ratings

The four evaluative ratings (liking, wanting, comfort, pleasantness) formed a highly internally consistent scale (see Supplementary Materials). Therefore, our outcomes were ‘touch evaluation’ (four ratings entered concurrently into multivariate linear mixed models; see *Plan of Analysis*) and touch intensity (single rating).

There was an effect of velocity on evaluative touch ratings that was not qualified by country or body region (see Table 2): participants in SA and UK evaluated affective, slow touch (*M* = 73.47, *SE* = 2.48) significantly more positively than the faster touch (*M* = 67.34, *SE* = 2.47), in line with our hypothesis, though the faster touch was still evaluated moderately positively. Ratings did not differ by body region. Examining effects of country, participants in SA evaluated the dynamic touch significantly more positively (*M =* 76.78, *SE =* 3.37) and significantly less intense (*M =* 23.63, *SE =* 5.00) than UK participants (evaluative rating: *M =* 61.54, *SE =* 4.08; intensity rating: *M =* 56.02, *SE =* 6.06) across stroking velocities and body regions (see Table 2), in line with our hypothesis.

**Table 2.**
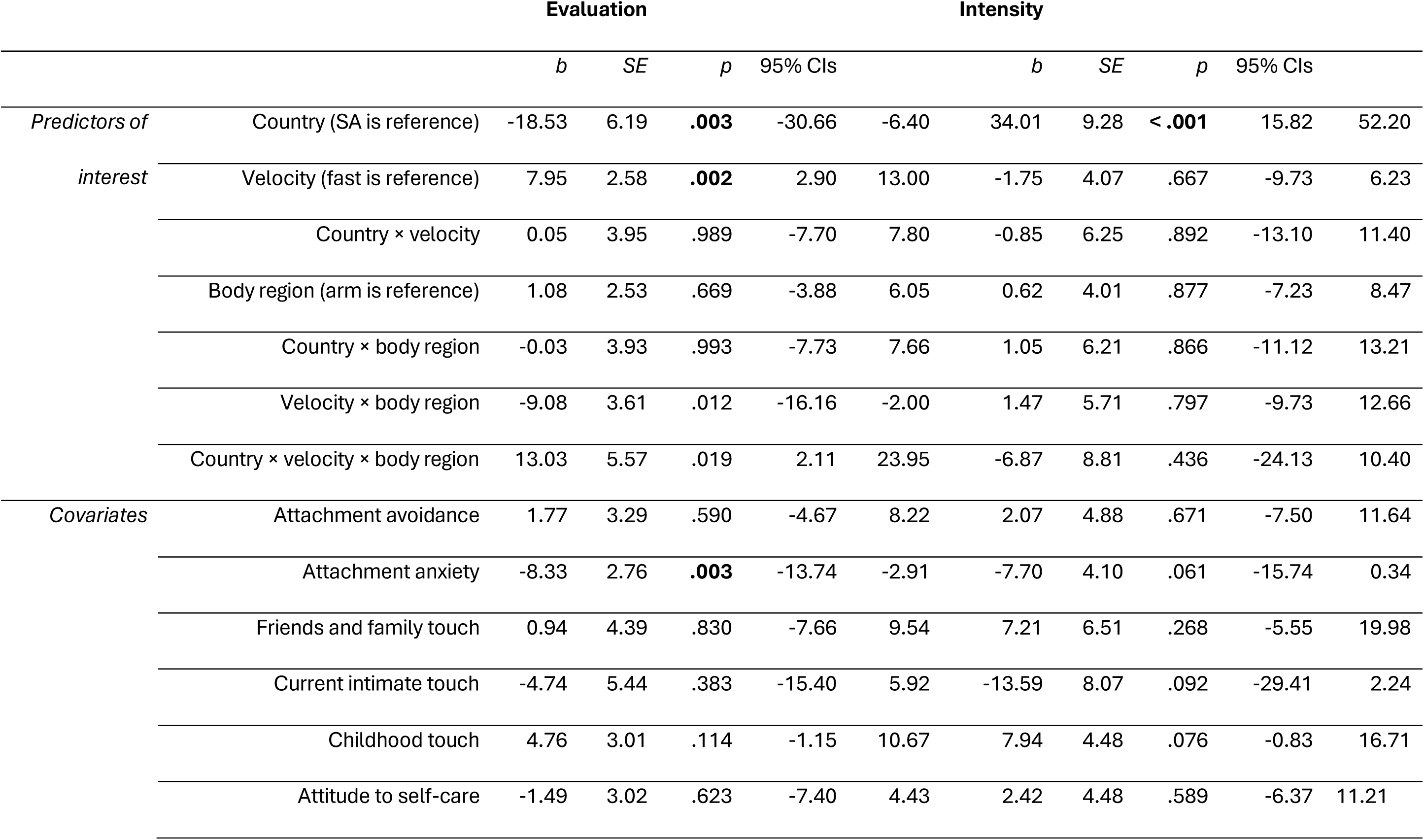
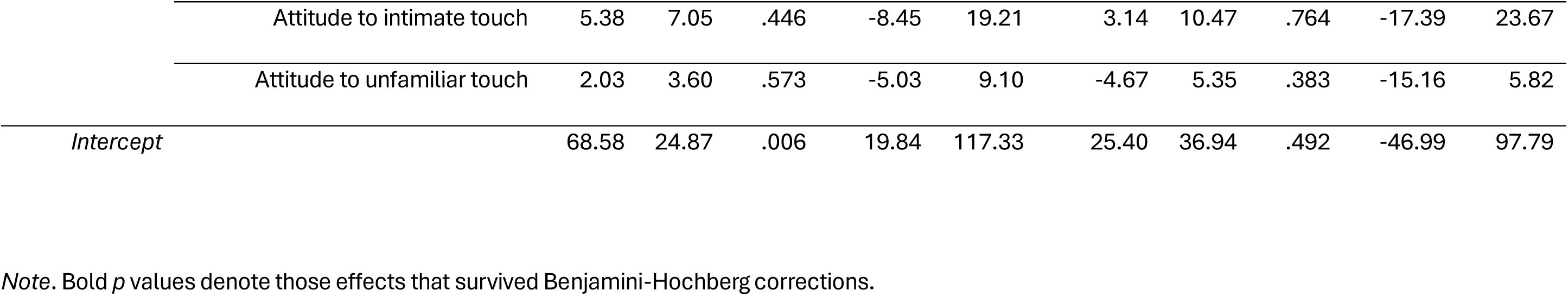
Linear mixed modelling results for evaluative (multivariate) and intensity (univariate) touch rating outcomes.

|  |  | Evaluation |  |  |  | Intensity |  |  |  |  |  |
| --- | --- | --- | --- | --- | --- | --- | --- | --- | --- | --- | --- |
|  |  | *b* | *SE* | *p* | 95% CIs |  | *b* | *SE* | *p* | 95% CIs |  |
| *Predictors of interest* | Country (SA is reference) | -18.53 | 6.19 | **.003** | -30.66 | -6.40 | 34.01 | 9.28 | **< .001** | 15.82 | 52.20 |
|  | Velocity (fast is reference) | 7.95 | 2.58 | **.002** | 2.90 | 13.00 | -1.75 | 4.07 | .667 | -9.73 | 6.23 |
|  | Country × velocity | 0.05 | 3.95 | .989 | -7.70 | 7.80 | -0.85 | 6.25 | .892 | -13.10 | 11.40 |
|  | Body region (arm is reference) | 1.08 | 2.53 | .669 | -3.88 | 6.05 | 0.62 | 4.01 | .877 | -7.23 | 8.47 |
|  | Country × body region | -0.03 | 3.93 | .993 | -7.73 | 7.66 | 1.05 | 6.21 | .866 | -11.12 | 13.21 |
|  | Velocity × body region | -9.08 | 3.61 | .012 | -16.16 | -2.00 | 1.47 | 5.71 | .797 | -9.73 | 12.66 |
|  | Country × velocity × body region | 13.03 | 5.57 | .019 | 2.11 | 23.95 | -6.87 | 8.81 | .436 | -24.13 | 10.40 |
| *Covariates* | Attachment avoidance | 1.77 | 3.29 | .590 | -4.67 | 8.22 | 2.07 | 4.88 | .671 | -7.50 | 11.64 |
|  | Attachment anxiety | -8.33 | 2.76 | **.003** | -13.74 | -2.91 | -7.70 | 4.10 | .061 | -15.74 | 0.34 |
|  | Friends and family touch | 0.94 | 4.39 | .830 | -7.66 | 9.54 | 7.21 | 6.51 | .268 | -5.55 | 19.98 |
|  | Current intimate touch | -4.74 | 5.44 | .383 | -15.40 | 5.92 | -13.59 | 8.07 | .092 | -29.41 | 2.24 |
|  | Childhood touch | 4.76 | 3.01 | .114 | -1.15 | 10.67 | 7.94 | 4.48 | .076 | -0.83 | 16.71 |
|  | Attitude to self-care | -1.49 | 3.02 | .623 | -7.40 | 4.43 | 2.42 | 4.48 | .589 | -6.37 | 11.21 |

|  |  |  |  |  |  |  |  |  |  |  |
| --- | --- | --- | --- | --- | --- | --- | --- | --- | --- | --- |
| Attitude to intimate touch | 5.38 | 7.05 | .446 | -8.45 | 19.21 | 3.14 | 10.47 | .764 | -17.39 | 23.67 |
| Attitude to unfamiliar touch | 2.03 | 3.60 | .573 | -5.03 | 9.10 | -4.67 | 5.35 | .383 | -15.16 | 5.82 |
| *Intercept* | 68.58 | 24.87 | .006 | 19.84 | 117.33 | 25.40 | 36.94 | .492 | -46.99 | 97.79 |

*Note.* Bold *p* values denote those effects that survived Benjamini-Hochberg corrections.

### Electroencephalography

We examined relative band power changes (ERD/ERS) occurring during touch separately for alpha (central and parietal electrode clusters), beta (central and parietal sites), and theta (frontal and central sites) frequency bands (see Supplementary Materials for full information). We first tested effects of touch velocity and body region across countries, and then included country (and interactions with touch velocity and body region) as fixed predictors of interest, as pre-registered. As we found that country interacted with velocity and body region, we present those analyses below (see Supplementary Materials for findings across countries).

### ERD/S findings

Grand-averaged ERD/S in alpha, beta and theta bands in each of the four conditions, averaged over all participants, are presented in Supplementary Figures 1 and 2. Cortical activation changes during dynamic touch as a function of velocity, body region, and country were evaluated using linear mixed effects models. Below, we present the effects by country that survived Benjamini-Hochberg correction in the text; see the tables for full results and Supplementary Materials for effects across countries.

#### Alpha band

No effects of country (alone or in interaction with velocity and/or body region) survived Benjamini-Hochberg corrections (see Table 3). However, there was an effect of velocity across countries at central sites. Alpha-band ERS was significantly greater for slow-velocity touch (*M =* -.32, *SE =* 3.43) compared to ERD at faster-velocity touch (*M =* 5.41, *SE =* 3.42) across body regions. There was also an effect of body region at both central and parietal alpha sites. Alpha-band ERS was significantly larger for the arm vs. greater ERD at palm at central (*M =* -.58, *SE =* 3.43 for arm; *M =* 5.81, *SE =* 3.43 for palm) and parietal sites (*M =* -8.99, *SE =* 1.72 for arm; *M =* -5.33, *SE =* 1.72 for palm) across stroking speeds. Therefore, alpha band oscillations were not influenced by cultural context but were modulated by stroking speed and body region.

**Table 3.**
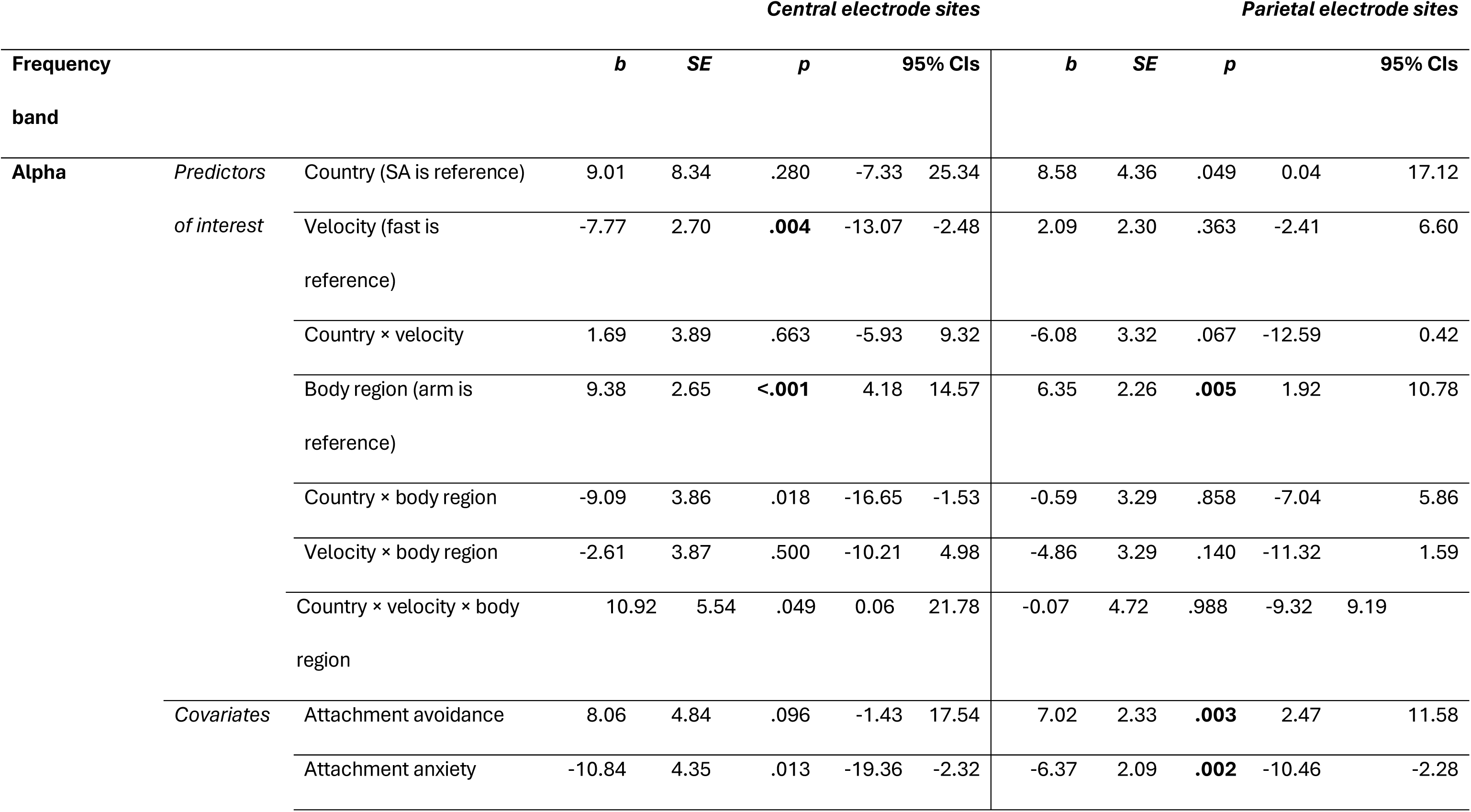
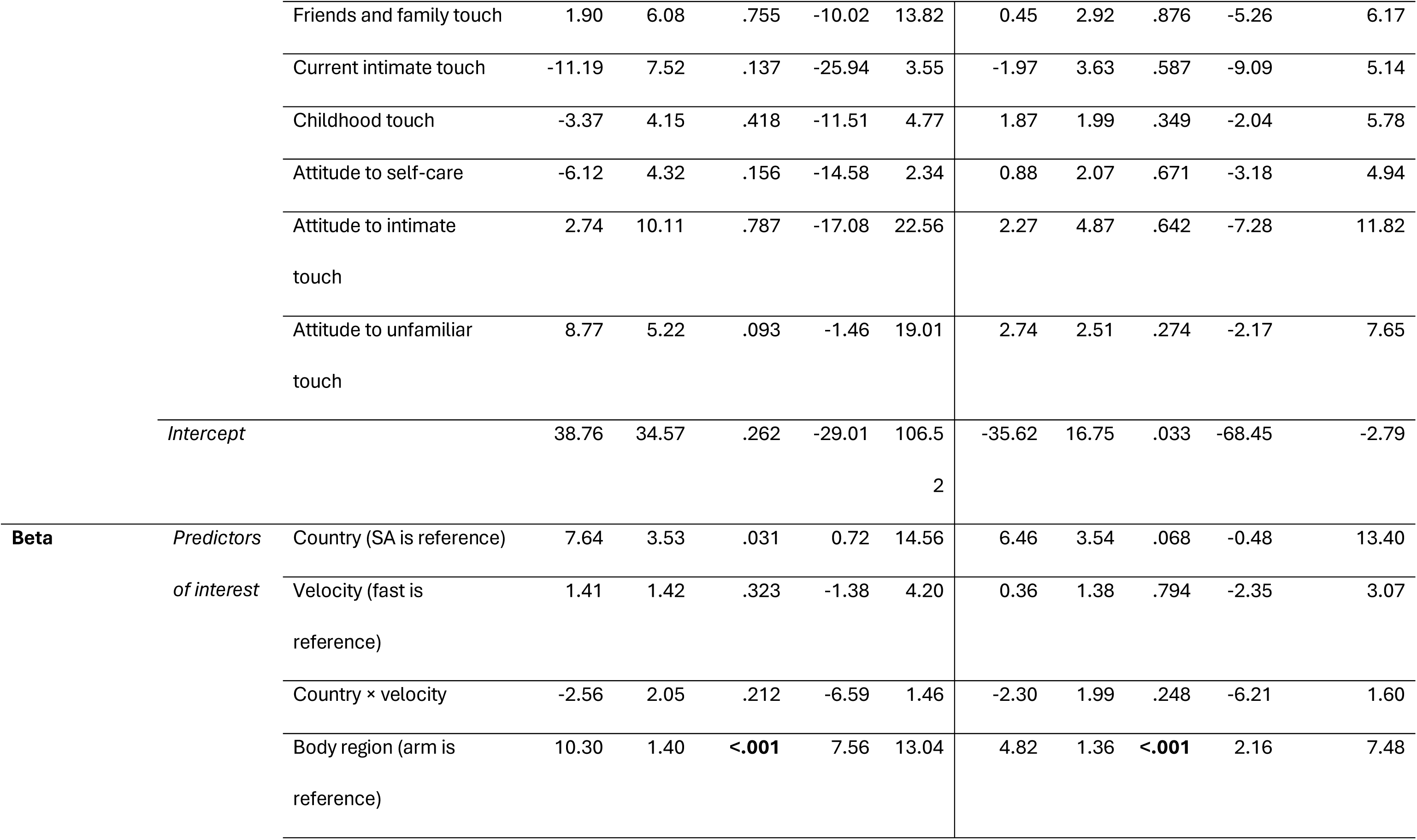
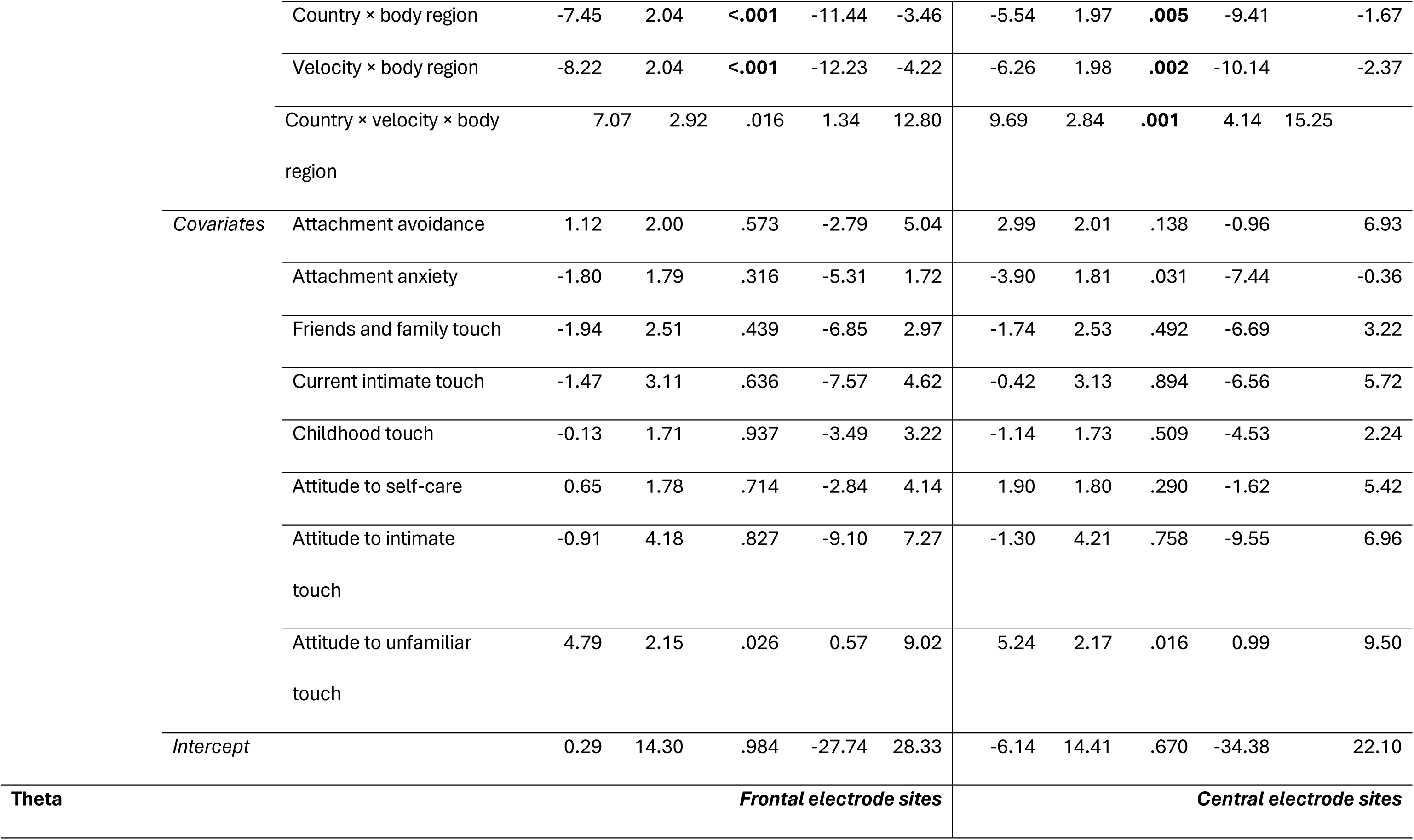
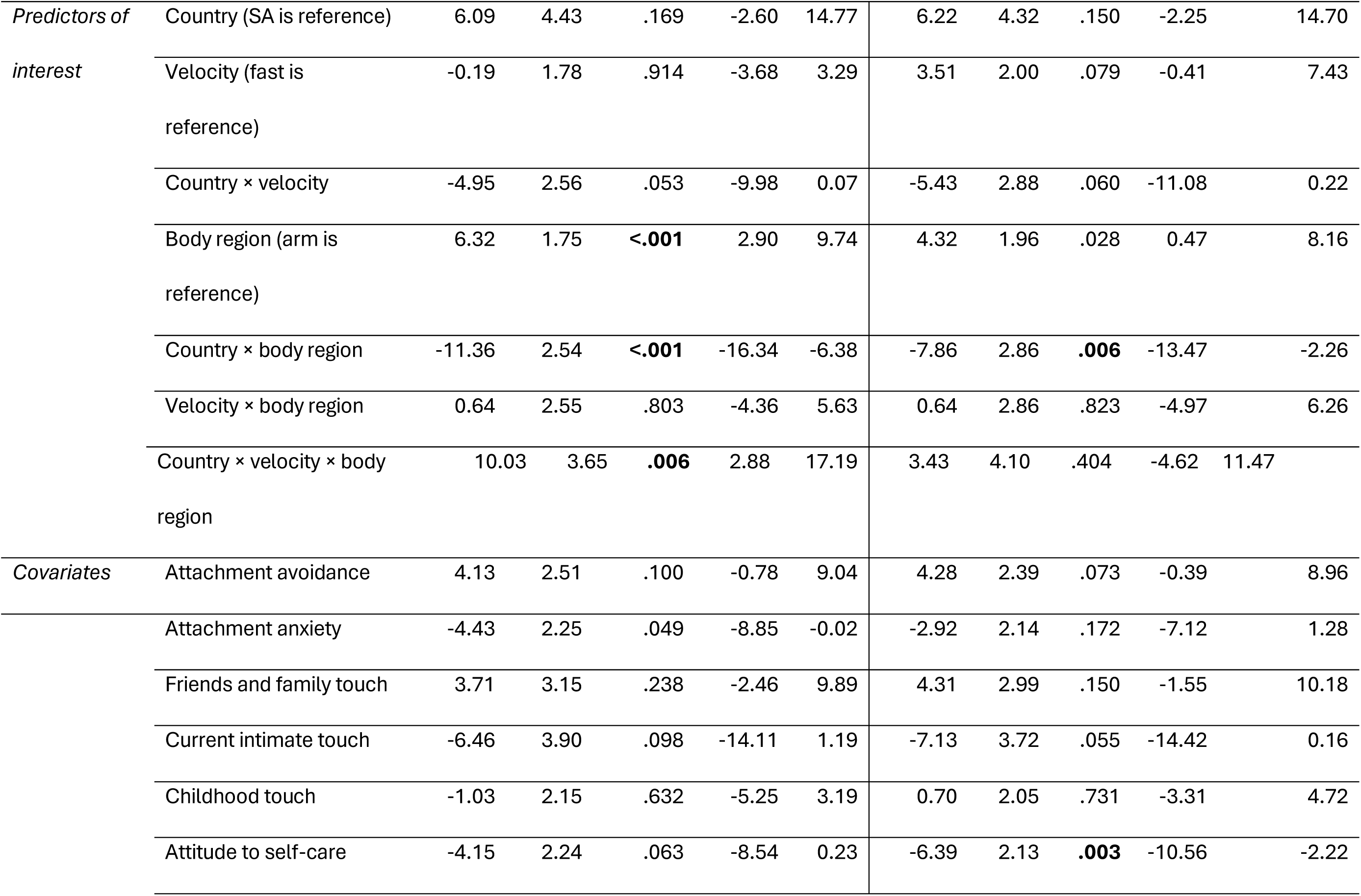
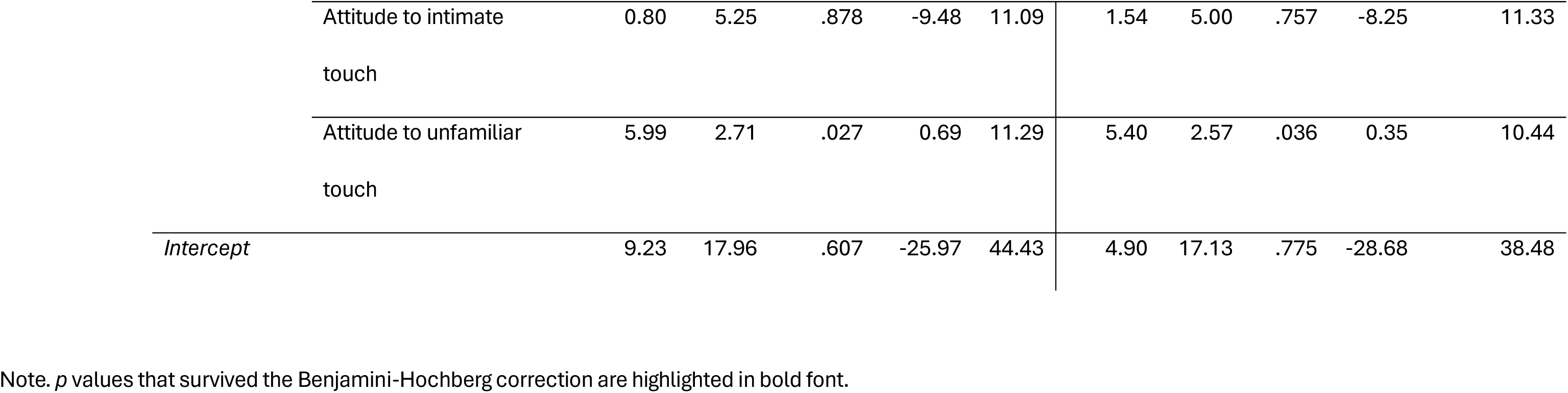
Multivariate linear mixed modelling results for country effects on alpha, beta, and theta ERD/S.

#### Beta band

For both central and parietal electrode sites, the effects of body region were shaped by cultural context (Figure 1b, e). The difference in the strength of beta-band ERD between palm and arm in bilateral central electrodes was significant in SA and UK participants, but was more pronounced in SA. SA participants showed larger ERD in central electrodes touch to the palm vs. ERS during touch to the arm than in UK participants (see planned contrast statistics in Figure2a, b). Planned contrasts were not significant for beta-band ERD in parietal electrodes, though the trend was in the same direction as the central site (Figure 2c, d). The effect in parietal electrodes was further qualified by a three-way interaction of country, velocity, and body region (see Table 3). Breaking this interaction down by country, the velocity by body region interaction for beta-band ERD/S in central electrodes was significant in SA (*b =* -8.27, *SE =* 2.24, *p =* .005) and UK participants (*b =* 3.44, *SE =* 1.68, *p =* .041); there was stronger ERS for slow vs. fast touch at the palm but not the arm in SA participants, whilst none of the planned contrasts were significant in the UK sample (Figure 2c, d). These results are in line with our hypothesis that SA (vs. UK) participants would show enhanced differentiation in their neural activation patterns.

**Figure 1.**
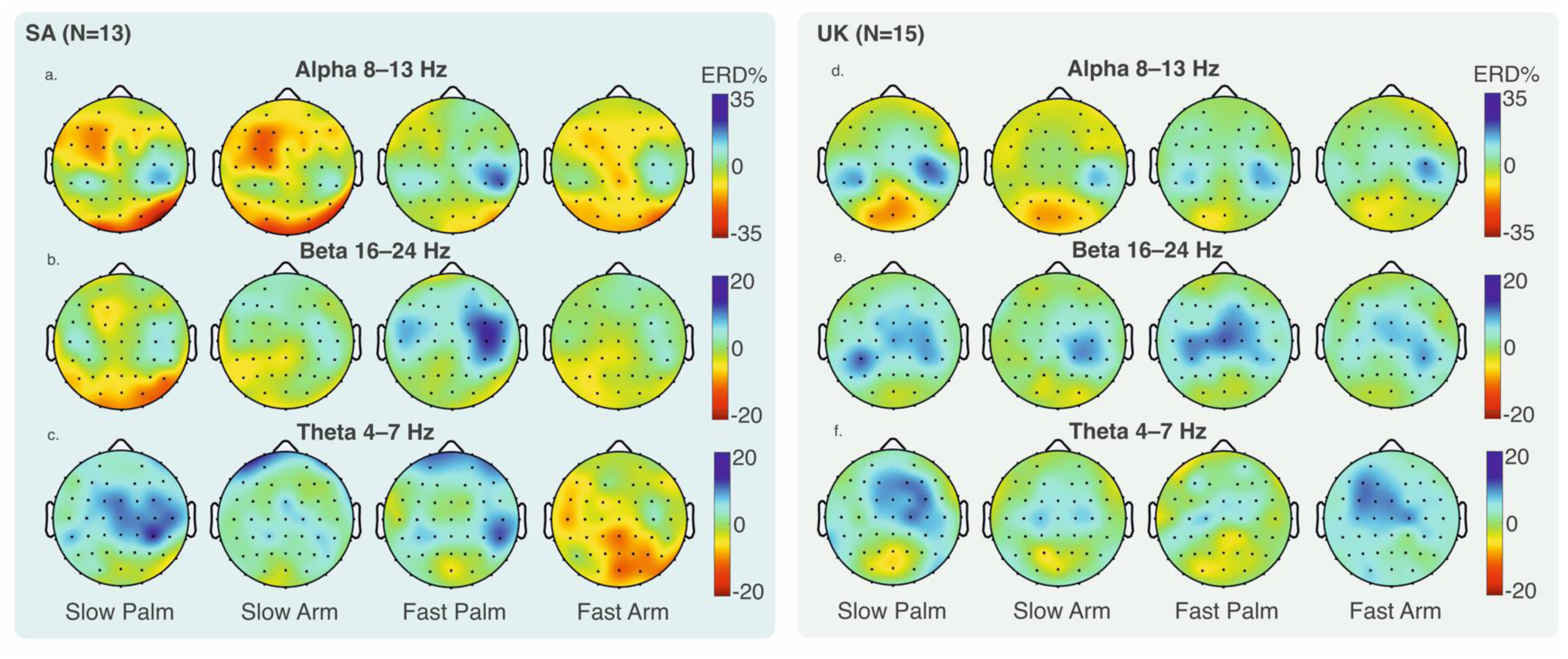
Grand-averaged event-related desynchronisation/synchronisation (ERD/S) during touch (0-3 s) by country in alpha (a, d), beta (b, e) and theta (c, f) bands in each of the four conditions, for SA and UK participants. Only participants with data for all conditions are shown in the topographic plots (as matrices need to be equal), resulting in a slightly smaller n here than the number included in statistical analyses. Topographic plots show only electrodes which were common across countries.

**Figure 2.**
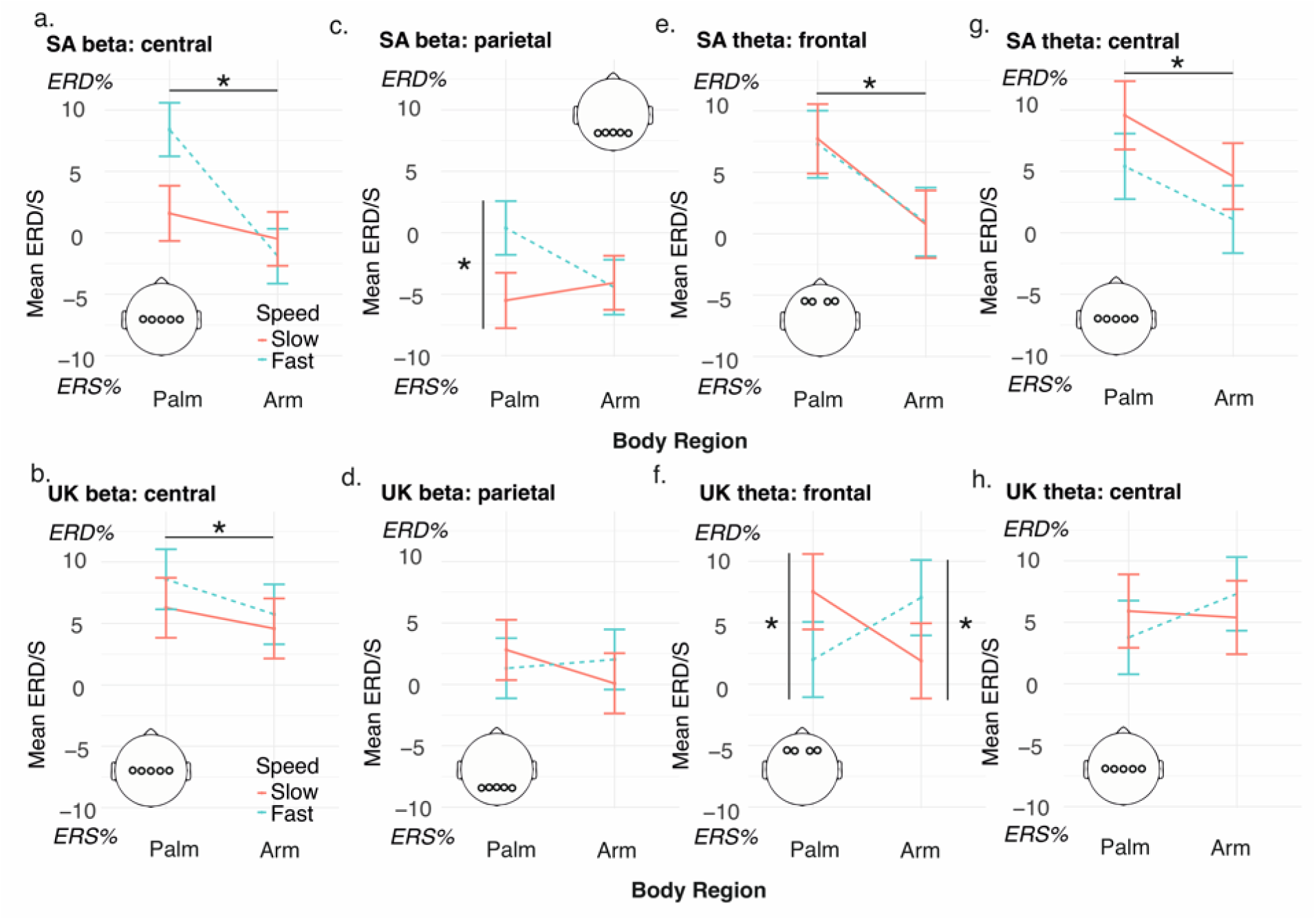
Effects of country plotted by body region (x axis) and velocity (separate lines; fast = dotted lines). Error bars denote ±1 standard error of the mean. For central beta, the difference between palm and arm (across velocities) was significant in the SA (a; Planned contrast palm vs. arm = 6.19, SE = 1.0, p < .001) and the UK sample (b; contrast = 2.27, SE = 1.05, p = .030). For parietal beta ERD/S in SA participants (c), the slow vs. fast contrast was significant only at the palm (contrast slow vs. fast = -5.92, SE = 1.57, p < .001) and not the arm (contrast slow vs. fast = .34, SE = 1.56, p = .827) body region, while in the UK (d), neither of these Bonferroni-corrected contrasts were significant (palm slow vs. fast contrast = 1.50, SE = 1.19, p = .208; arm contrast = -1.94, SE = 1.19, p = .102). For frontal theta, ERD was reduced for the arm compared to palm body regions in SA participants (e; frontal theta contrast = 6.64, SE = 1.24, p < .001) but not UK participants (f; frontal theta contrast = .30, SE = 1.31, p = .861). However, in the UK sample (f), the slow vs. fast contrast was significant at both palm (contrast = 5.52, SE = 1.75, p = .002) and arm (contrast = -5.15, SE = 1.75, p = .003) body regions, but in opposite directions, with greater ERD for slow vs. fast touch at the palm, and lower ERD for slow vs. fast touch at the arm body region. For central theta, ERD was reduced for the arm compared to palm body regions in SA participants (panel g; central theta contrast = 4.64, SE = 1.40, p = .001) but not UK participants (panel h; central theta contrast = -1.51, SE = 1.47, p = .303).

#### Theta band

There was a country by body region interaction in theta-band ERD/S in both frontal and central electrode sites (Table 3; Figure 2e-h). At both sites, as for beta-band oscillations, theta-band ERD was reduced for the arm compared to palm region in SA participants (Figure 2e, g) but not UK participants (Figure 2f, h), pointing to greater differentiation between body regions but not stroking speeds in SA participants. However, the 3-way interaction of country by velocity by body region was also significant for theta-band in frontal electrodes. Here, the velocity by body region interaction was significant in the UK (*b* = 10.67, *SE =* 2.48, *p <* .001) but not the SA sample (*b =* .60, *SE* = 2.66, *p* = .820). In the UK sample, the slow vs. fast contrast was significant at both palm and arm body regions (Figure 2f), but in opposite directions, with larger ERD for slow vs. fast touch at the palm, and lower ERD for slow vs. fast touch at the arm. While SA participants differentiated between body regions, UK participants additionally differentiated between stroking speeds, with opposite directions of effects at arm and palm.

Taken together, effects of body region (arm versus palm) on ERD/S were more pronounced in the SA sample, with larger beta- and theta-band ERD for the palm, and larger ERS for slow (vs. fast touch) also only evident at the palm body region for SA participants. In the UK, by contrast, effects of body region and stroking velocity were evident in theta-band oscillations at frontal sites, with larger ERD for slow touch on the palm, and reduced ERD for slow touch to the arm. Alpha ERD did not appear to be modulated by cultural context. Instead, alpha-band ERD in central electrodes was generally reduced for slow vs. fast touch, and relative band power was decreased (attenuated ERD and greater ERS) for the arm vs. palm at central and parietal sites.

## Discussion

This is the first study to experimentally investigate the influence of cultural context on self-reported and neurophysiological responses to affective (slow, dynamic) touch, accounting for individual differences in touch experiences and attitudes. We compared women living in SA, an underrepresented majority-world context ^12^, with women living in the UK, in a split-site, laboratory-based study. We found that cultural context modulated both the affective (how pleasant, comfortable, liked, and wanted the touch was rated to be) and intensity evaluations of dynamic stroking touch, as well as cortical oscillatory patterns (ERD/S) in beta and theta bands. Evaluation of stroking speed and ERD/S in alpha band did not differ between countries.

Participants in SA rated all types of touch – irrespective of stroking speed or body region – more positively and as less intense than UK participants, in line with our hypothesis. Prevalence of affective touch is higher ^13^ and touch norms more positive ^15^ in countries and contexts that share characteristics with SA, but the perception of directly-experienced affective touch in people living in SA has not been explored. We found that dynamic touch is indeed evaluated in more positive terms by adults living in SA (compared to in the UK). We also observed that, across countries, slow-velocity touch was evaluated more positively than faster touch (replicating a robust main effect in the literature ^20,40^). This effect was evident across both body regions (cf. ^41^), supporting findings that top-down modulatory factors beyond activation of CT-afferents contribute to the sensation and evaluation of affective touch ^42^. We also asked participants about their general attitudes towards touch (beyond dynamic stroking) from intimate and unfamiliar others, and about current levels of intimate touch, friends and family touch, levels of positive childhood touch, and adult attachment styles. No country differences were observed on these measures, except for attitudes to unfamiliar touch, which were less positive in SA (vs. UK) participants. The discrepancy in touch evaluation vs. touch attitudes may be explained by the study environment, in which tactile stimulation was delivered in a controlled, arguably safe way, by a trained experimenter of the same gender as participants. By contrast, attitudes to unfamiliar touch may be more generally influenced by high rates of interpersonal violence in SA ^43^.

Examining amplitude changes of cortical oscillations in response to slow- and faster-velocity touch at arm and palm body regions, we observed modulation by cultural context in beta and theta bands, and effects across countries in alpha band. Participants in SA showed enhanced differentiation between touch to body regions in the beta band. Specifically, participants in SA showed larger ERD over central scalp regions during touch to the palm (vs. ERS to the arm) than UK participants, and stronger parietal ERS for slow vs. fast touch at the palm (but not the arm); a contrast that was not significant in the UK sample. Sensorimotor beta oscillations are proposed to play an active role in endogenous top-down processing and sensorimotor integration by linking sensory input with contextual knowledge ^44^. Tactile expectations have been shown to modulate pre-stimulus beta oscillations in the primary somatosensory cortex, with this effect enhanced by attention ^45^. Studies in primates show that sensorimotor beta oscillations correspond to somatosensory decision outcomes and are modulated only by stimulus features when task-relevant ^46^; a finding also supported by human EEG studies ^47^. Thus, enhanced desynchronisation of central beta oscillations during touch may represent context-specific endogenous modulation shaping the neural processing of touch, particularly in during fast touch and touch to the palm.

Heightened cortical activation signified by ERD might also be linked to the palm’s greater innervation density and smaller receptive fields ^48^ and greater tactile acuity ^49^ when compared to the forearm. These features facilitate precise sensory processing, especially during tasks requiring detailed mapping of stimulus characteristics, such as object manipulation and grasping ^50^. The hands have larger cortical somatotopic representations compared to the forearms and much of the rest of the body, further emphasising their importance in detailed sensory processing ^51^. However, since innervation density is unlikely to differ across cultures, our findings suggest that top-down modulatory processes, rather than sensory differences, influenced sensory processing in SA participants.

The significance of the palm as a communication channel may extend beyond its sensory attributes to cultural and social dimensions. Previous research has found differences in touch permissibility to different body sites as a function of emotional closeness; for example, stranger touch is more acceptable to the hand than the forearm ^52^, and cultural differences in terms of the strength of this association for arm vs. hand in British vs. Japanese cultural contexts ^16^. Moreover, Chinese participants preferred touch to the hands more than German participants ^14^. The arm and hand may therefore hold different significance in different cultural contexts, and further research is needed to investigate our effect of heightened cortical engagement in relation to palm vs. arm touch in participants living in SA. Handshakes play an important role in traditional greetings across Africa (see ^53^). In SA, palm touch, as involved in the three-point handshake, may hold augmented social significance, but this remains to be fully explored.

Regarding theta-band oscillations, previous affective and social touch studies found stronger theta-band ERD in response to slow, CT-optimal touch (compared to faster non-CT-optimal touch ^32^) and hand-holding ^33^, suggesting a potentially soothing effect of pleasant and prosocial touch. In SA participants, we did not find differentiation between slow and faster touch, but, across touch speeds, theta-band ERD over central and frontal sites was reduced for the arm compared to palm region in SA participants. While we are unaware of studies that have contrasted theta-band oscillations during dynamic touch across the arm and palm, this finding might again point to a greater significance of palm touch in SA participants. In contrast, UK participants differentiated between touch velocities at each body region, with greater ERD for slow vs. fast touch at the palm, and lower ERD for slow vs. fast touch at the arm. This finding does not support von Mohr, et al. ^32^, though their faster touch velocity was 32cms^−1^, compared to 18cms ^−1^ in the current study. As increased frontal theta-band power is linked with somatosensory orienting ^54^ as well as attention and cognitive control ^30^, slow palm stroking may also be more unusual and therefore perceptually salient, though this remains to be empirically tested. Faster stroking touch has also been associated with communicating intentions of warning ^3^, potentially also explaining heightened attention during faster stroking speeds. Collectively, these effects likely reflect differences in the social or emotional importance of touch in each cultural context.

Alpha band power was not modulated by cultural context, but we found general effects across participants of both countries. Specifically, faster touch (over both body regions) induced stronger central alpha ERD, and touch applied to the palm region was also associated with greater central alpha ERD and reduced parietal alpha ERS compared to the arm. 10 Hz and 20 Hz ERD in somatosensory regions have been associated with cortical excitability and readiness for sensory processing ^55^, and ERD amplitude reflecting attentional resource allocation and arousal ^56^. These processes are further shaped by social factors, with increased alpha power over frontal and parietal regions found when participants were resting (not engaging in an explicit task) with somebody else present vs. resting alone ^57^. Moreover, Verbeke, et al. ^57^ found that greater attachment anxiety was associated with increased frontal and parietal alpha only in the together condition. Given we only had a ‘together’ condition in this study, it is interesting to note a positive link between attachment anxiety and avoidance and parietal alpha-band ERD in our results. In sum, alpha-band oscillations were not modulated by cultural context, indicating that while potentially shaped by individual differences, these attentional processes may not be culturally specific.

A strength of our study is investigating not only cross-cultural differences but also measuring individual differences relating to touch attitudes and experiences and controlling for inter-individual variance in analyses. Thus, our findings are not due to e.g., differences in individual attachment styles or current intimate touch levels across the two countries. By examining neural oscillatory patterns in response to slow vs. fast touch at both the arm and palm, we have also extended previous studies by contrasting touch to CT-innervated, hairy skin with touch to glabrous (sparsely or non-CT-innervated) skin. Moreover, we asked participants to rate touch in relation to five aspects (pleasantness, comfort, intensity, liking, and wanting) and found that the four affective terms formed an internally consistent scale. In future, researchers may choose to include one or more of these aspects that seem to relate to similar evaluative constructs, at least at the self-report level.

Several limitations should be noted. The samples used in this study are not representative of the entire population living in SA and the UK, nor were all our participants South African or British nationals. The term ‘culture’ is employed here as a wide generalisation to encompass South African compared to British social contexts. However, both SA and the UK are home to individuals from various countries and ethnic backgrounds, which likely influence their touch norms and experiences. Future studies would benefit from looking at more specific cultural groups within these countries to investigate more nuanced cross-cultural differences in affective touch experiences. Nevertheless, experimental studies that draw on neuroimaging methods (i.e., EEG) in the general field of cognitive neuroscience are rare and challenging to conduct in contexts such as South Africa (see ^58,59^), resulting in further underrepresentation of populations from Africa in EEG and experimental neuroscience research ^60^. Therefore, despite the above limitations, the current study is a critical step in adding to the diversity and representation of majority-world participants, specifically in Africa, in neuroimaging and experimental studies in cognitive neuroscience in general and in the affective touch literature more specifically.

It is also important to acknowledge that early touch experiences might be different for people who moved to SA or UK in adulthood, and touch norms can differ based on heritage for people now living in the same cultural context (see e.g., ^15^). Moreover, although we characterised our samples in terms of demographic and individual differences variables, we did not account for variables such as socioeconomic status. Regarding our individual difference measures, the touch experiences and attitudes measure has recently been validated in a large cohort of people living in South Africa ^61^. However, attachment style is a Western construct and may not apply universally ^62^. We did not find differences on our attachment style measure between SA and UK participants, but it may not be the most appropriate way of assessing mental representations of social relationships across cultural contexts.

In conclusion, our study provides the first experimental and neuroimaging evidence showing that cultural context modulates both subjective and neurophysiological responses to dynamic, affective touch. Considering subjective touch evaluation, we observed main effects of cultural context across body region and velocity. The neural data paints a more nuanced picture, suggesting there may be a disconnect between cortical representations and subjective touch evaluations. Taken together, extrapolating findings from a given study to populations in a different cultural context is unlikely to be appropriate, highlighting the need to consider top-down influences at every stage of touch processing. We hope this study will engender more cross-cultural touch research – especially in underrepresented majority-world contexts – into touch perception and evaluation, and touch norms, to allow us to gain a fuller picture of how touch is experienced and processed in different contexts. Moreover, we strongly recommend that researchers developing and implementing touch-based interventions (cf. ^39^) consider such country-level factors as well as individual differences in their design, and in the interpretation of their findings.

## Methods

The study design and methods were pre-registered on the Open Science Framework: https://osf.io/fcqnk. Ethical approval for the study was granted by the Institute of Population Health Research Ethics Committee, University of Liverpool, and the Human Research Ethics Committee (Medical) at the University of the Witwatersrand. Data collection ran from June – August 2022 in the UK and November 2023 – January 2024 in SA.

### Design and Procedure

The study employed a 2 (country i.e., UK, SA; between-subjects factor) × 2 (touch stroking velocity i.e., slow, CT-optimal 3cms ^−1^ vs. faster non-CT-optimal 18cms ^−1^; within-subjects factor) × 2 (body region i.e., CT-innervated forearm vs. non-CT-innervated palm; within-subjects factor) mixed design. Participants in both countries received slower and faster velocity touch to the arm and palm of the hand, with speed and body region counterbalanced across participants, while their EEG was recorded.

Outcome measures were: (1) self-reported pleasantness, comfort, intensity, liking and wanting ratings of touch; and (2) neural oscillations in theta, alpha, and beta frequency bands. We also measured self-reported experiences and attitudes to touch ^63^ and adult attachment style ^64^.

Volunteers were invited to participate in a study exploring brain activity associated with tactile pleasantness and touch velocity. Participants were requested to avoid caffeine intake on the day of the experiment and to arrive with clean, dry hair free from conditioner or hair products. Lab layouts are shown in Figure 3. In both testing sites, upon obtaining written informed consent, participants were seated in an upright position and asked to rest their right arm on the table behind a screen, with their left palm facing upwards. The screen was introduced to prevent the participant from seeing the experimenter or the touch during tactile stimulation. Participants completed questionnaires using the electronic platform Qualtrics while the EEG cap was applied. Thereafter, touch was administered while EEG data was recorded. After all blocks were completed, the EEG cap was removed, and participants were fully debriefed and compensated for their time.

**Figure 3.**
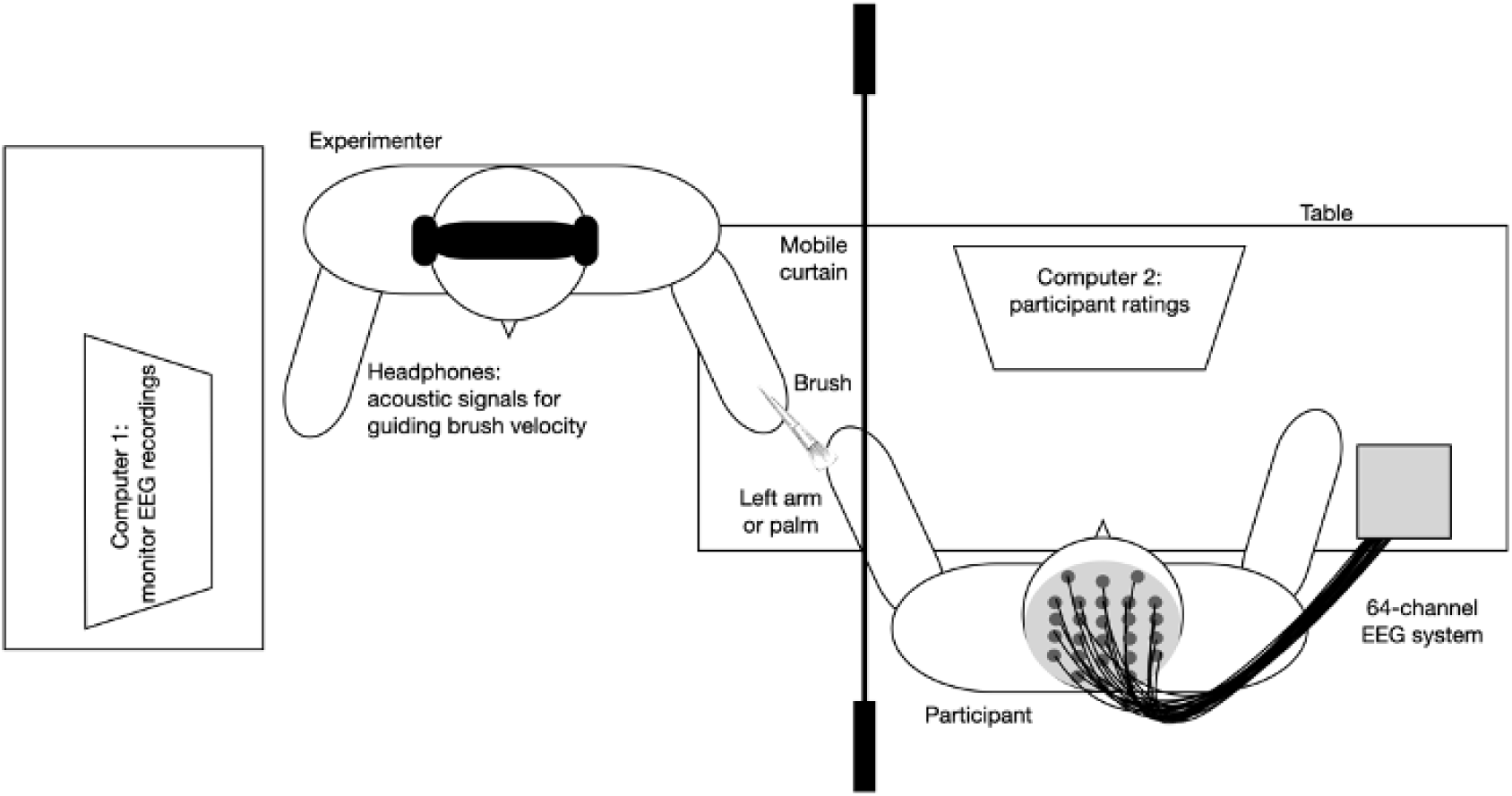
The schema depicts the experimental set-up in South Africa and in the United Kingdom. Slight alterations in the room set-up were made due to constraints to the physical laboratory space but the main layout and materials used were standardised and consistent across both study sites.

### Participants

*N* = 36 female participants were recruited in two different countries. *N* = 15 participants were recruited and took part in the study in Liverpool, UK, and *N* = 21 participants in Johannesburg, SA. The sample size was based on previous research (*N* = 28 in the similar EEG study by ^32^), though we included the exploratory country and individual differences variables, and we exceeded our target of *N* = 30 indicated in the OSF pre-registration.

We recruited only female participants, given biological sex differences in touch perception ^43^. All participants reported their gender identity as female. All experimenters were female to eliminate any possible confounding effects of gender on the social modulation of touch experiences. Participants were all aged 18 or over (*M =* 23.14 years, *SD =* 7.32 in SA, and *M =* 25.93 years, *SD =* 6.39 in UK) and there was no significant difference in terms of age between countries (see Table 4 for full demographic characteristics and difference tests). We specified that all participants must be right-handed, as touch was administered to the non-dominant left arm and palm. Exclusion criteria were current or history of psychiatric disorder, or neurological or medical conditions, which might influence touch perception (e.g., chronic pain and associated allodynia), and having wounds, scars, tattoos or skin irritation/diseases on their forearms where touch was administered.

**Table 4.**
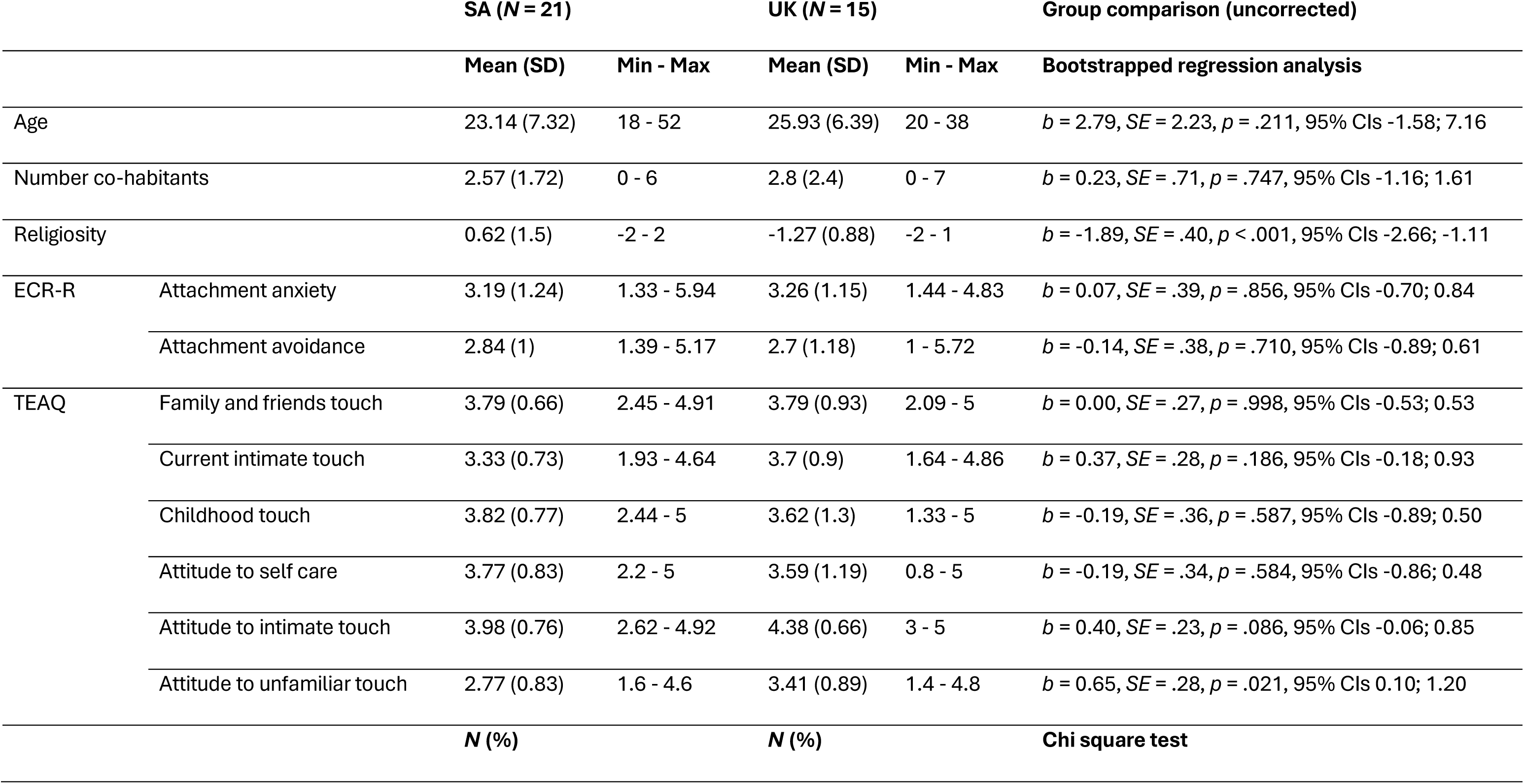
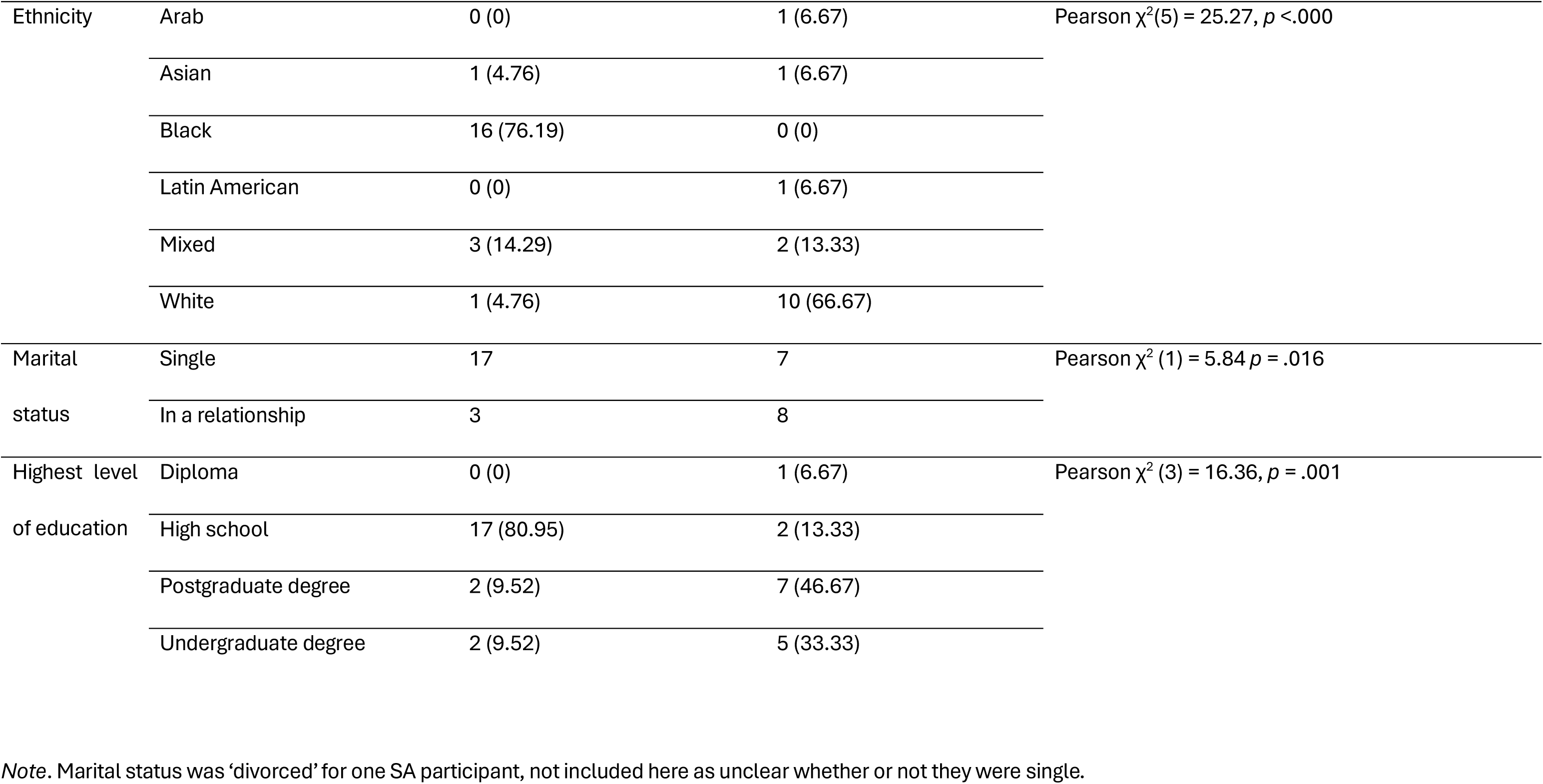
Demographic characteristics and self-report questionnaires.

|  |  | SA (N = 21) |  | UK (N = 15) |  | Group comparison (uncorrected) |
| --- | --- | --- | --- | --- | --- | --- |
|  |  | Mean (SD) | Min - Max | Mean (SD) | Min - Max | Bootstrapped regression analysis |
| Age | | 23.14 (7.32) | 18 - 52 | 25.93 (6.39) | 20 - 38 | $b = 2.79, SE = 2.23, p = .211, 95\% \text{ CIs } -1.58; 7.16$ |
| Number co-habitants | | 2.57 (1.72) | 0 - 6 | 2.8 (2.4) | 0 - 7 | $b = 0.23, SE = .71, p = .747, 95\% \text{ CIs } -1.16; 1.61$ |
| Religiosity | | 0.62 (1.5) | -2 - 2 | -1.27 (0.88) | -2 - 1 | $b = -1.89, SE = .40, p < .001, 95\% \text{ CIs } -2.66; -1.11$ |
| ECR-R | Attachment anxiety | 3.19 (1.24) | 1.33 - 5.94 | 3.26 (1.15) | 1.44 - 4.83 | $b = 0.07, SE = .39, p = .856, 95\% \text{ CIs } -0.70; 0.84$ |
| | Attachment avoidance | 2.84 (1) | 1.39 - 5.17 | 2.7 (1.18) | 1 - 5.72 | $b = -0.14, SE = .38, p = .710, 95\% \text{ CIs } -0.89; 0.61$ |
| TEAQ | Family and friends touch | 3.79 (0.66) | 2.45 - 4.91 | 3.79 (0.93) | 2.09 - 5 | $b = 0.00, SE = .27, p = .998, 95\% \text{ CIs } -0.53; 0.53$ |
| | Current intimate touch | 3.33 (0.73) | 1.93 - 4.64 | 3.7 (0.9) | 1.64 - 4.86 | $b = 0.37, SE = .28, p = .186, 95\% \text{ CIs } -0.18; 0.93$ |
| | Childhood touch | 3.82 (0.77) | 2.44 - 5 | 3.62 (1.3) | 1.33 - 5 | $b = -0.19, SE = .36, p = .587, 95\% \text{ CIs } -0.89; 0.50$ |
| | Attitude to self care | 3.77 (0.83) | 2.2 - 5 | 3.59 (1.19) | 0.8 - 5 | $b = -0.19, SE = .34, p = .584, 95\% \text{ CIs } -0.86; 0.48$ |
| | Attitude to intimate touch | 3.98 (0.76) | 2.62 - 4.92 | 4.38 (0.66) | 3 - 5 | $b = 0.40, SE = .23, p = .086, 95\% \text{ CIs } -0.06; 0.85$ |
| | Attitude to unfamiliar touch | 2.77 (0.83) | 1.6 - 4.6 | 3.41 (0.89) | 1.4 - 4.8 | $b = 0.65, SE = .28, p = .021, 95\% \text{ CIs } 0.10; 1.20$ |
|  |  | N (%) |  | N (%) |  | Chi square test |

|  |  |  |  |  |
| --- | --- | --- | --- | --- |
| Ethnicity | Arab | 0 (0) | 1 (6.67) | Pearson $\chi^2(5) = 25.27, p < .000$ |
|  | Asian | 1 (4.76) | 1 (6.67) |  |
|  | Black | 16 (76.19) | 0 (0) |  |
|  | Latin American | 0 (0) | 1 (6.67) |  |
|  | Mixed | 3 (14.29) | 2 (13.33) |  |
|  | White | 1 (4.76) | 10 (66.67) |  |
| Marital status | Single | 17 | 7 | Pearson $\chi^2(1) = 5.84 p = .016$ |
|  | In a relationship | 3 | 8 |  |
| Highest level of education | Diploma | 0 (0) | 1 (6.67) | Pearson $\chi^2(3) = 16.36, p = .001$ |
|  | High school | 17 (80.95) | 2 (13.33) |  |
|  | Postgraduate degree | 2 (9.52) | 7 (46.67) |  |
|  | Undergraduate degree | 2 (9.52) | 5 (33.33) |  |

*Note.* Marital status was 'divorced' for one SA participant, not included here as unclear whether or not they were single.

On average, participants lived with two to three other people, and there was no significant difference between countries regarding number of co-habitants; however, SA participants were predominantly single, while half of UK participants were in a relationship. There was a significant difference in terms of ethnicity, with most SA participants identifying as black and most UK participants as white. Furthermore, there was a significant difference in highest level of education as most SA participants were undergraduate students, while more UK participants were postgraduate students. We assessed religiosity as a country-level variable previously associated with attitudes towards touch ^13^. SA participants were significantly more religious (we did not assess which religion) compared to UK participants.

### Materials and Measures

#### Tactile stimulation

Tactile stimulation was administered by a trained experimenter, unknown to the participants, using a cosmetic make-up brush (Natural Hair Blush Brush, No. 7, The Boots Company). Four 9 cm long × 4 cm wide areas were marked on the participant’s skin: two regions were marked contiguously along participants’ left volar forearm between wrist and elbow, and two areas were marked side-by-side on the surface of the palm. Touch was administered in a block design. Each block included 5 practice trials (not included in analyses) and 40 3-second trials. Trials were separated by 8-s of no touch, and adjacent skin areas were alternated between strokes, to prevent habituation and to allow cortical oscillatory band power to return to baseline.

Tactile stimulation was applied at two different velocities: a slower speed of 3cms ^−1^ which optimally activates C tactile afferent fibres, and a faster speed of 18cms ^−1^ which does not optimally activate these fibres (as in ^65,66^). Thus, there were four blocks (CT-innervated arm at 3cms ^−1^, CT-innervated arm at 18cms ^−1^, non-CT-innervated palm at 3cms ^−1^, non-CT-innervated palm at 18cms ^−1^). Each of the experimenters were trained in the touch protocol using a standardised video to ensure consistency in the administration of tactile stimulation across testing sites, and touch onsets were standardised with a countdown of 4 s beeps, which were audible to the experimenter through headphones. A virtual pilot session was also held with each site to observe and ensure constancy of the experimental and EEG protocol.

Participants rated pleasantness, comfort, intensity, liking and wanting ratings of touch after each block on visual analogue scales with the anchors 0 (*not at all*) to 100 (*extremely*).

#### Electroencephalography

EEG data was recorded from 64 active silver-silver chloride electrodes using a BrainProducts actiCap snap system (BrainProducts GmbH, Munich, Germany) for the UK study, and a g.tec g.Hlamp (g.tec medical engineering GmbH, Schiedllberg, Austria) for the SA study. For both studies, electrodes were embedded inside the electrode cap, which was securely positioned on the participant’s scalp, aligned with respect to three anatomical landmarks of two preauricular points and the nasion following the International 10-20 system. The BrainProducts actiCap system utilised Fz as the reference electrode and FPz as the ground electrode (resulting in 63 active recording electrodes). The g.tec system utilised linked earlobe references A1 and A2 and AFz as the ground electrode (resulting in 62 active recording electrodes). Electrolyte gel was applied to achieve sufficiently low electrode-to-skin impedances at the beginning of the experiment (<5kΩ for g.tec and <25kΩ for BrainProducts, as per manufacturer recommendations). Signals were digitised at 1kHz using an actiChamp (UK) or g.Hlamp (SA) DC amplifier. Data were stored directly onto the hard drive for subsequent offline processing and analysis.

#### EEG Data Processing

EEG data were processed using EEGLab ^67^. Continuous data were split into 8 s epochs (-2.5 to 5.5 s around touch onset) and combined into one datafile for each participant. Data were re-referenced to the common average, and filtered using 1 Hz high pass and 70 Hz low pass filters. A notch filter from 48–52 Hz was applied to remove mains line noise, and data were downsampled to 256 Hz.

Artefacts were removed using a semi-automated method. Oculomotor artefacts were removed from the data using independent component analysis. Next, electrode channels with large artefacts were identified with visual inspection and interpolated (maximum of 10% of total electrodes). Outlier values were identified using *pop_eegthresh.m* with limits of -125 – 125 µV. Improbable data were marked using *pop_jointprob.m* using single channel and global channel limits of 5 standard deviations. For each participant, data were visually inspected and marked trials were manually reviewed. Mean number of epochs remaining after artefact correction for each condition were: slow palm, 34.91 ± 5.93; fast palm, 36.15 ± 2.45; slow arm, 35.91 ± 3.32; fast arm, 35.59 ± 4.64. A univariate ANOVA with within-subject factors of body region and velocity and a between-subjects factor of country showed that there was no significant difference in accepted trials between touch blocks (*p* > .05). Significantly more trials were rejected from the SA (35.71 ± 3.01) compared to the UK (37.18 ± 2.22) dataset (*F*(1,127) = 9.04, *p =* .003).

Power spectra were computed in FieldTrip (http://fieldtriptoolbox.org) using a discrete Fourier time-frequency transformation. Power spectral densities were computed using Welch’s method from 1-s overlapping segments. Data were smoothed using a 4 Hz Slepian sequence prior to Fourier transformation. The spectral window was shifted in 0.1 second intervals to yield a power time series of 80 points. Spectral power was estimated in the range 1–70 Hz with a frequency resolution of 1 Hz. Relative power was evaluated using the classical ERD transformation (Pfurtscheller & Aranibar, 1979): 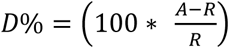 where D represents the percentage power change during epochs following stimulus onset (A) relative to baseline (R). Positive D values correspond to relative power decreases (event-related desynchronisation, ERD), while negative D values correspond to power increases (event-related synchronisation, ERS ^27,28^).

#### Questionnaires

Participants completed a series of questionnaires including demographic characteristics, the *Experiences in Close Relationships Revised Questionnaire* (ECR-R ^64^), and the *Touch Experiences and Attitudes Questionnaire* (TEAQ ^63^).

The ECR-R is a widely used assessment tool that measures individual differences in attachment-related anxiety and avoidance in adult romantic relationships. It consists of 36 items, including 18 items that assess attachment anxiety (‘I’m afraid that I will lose my partner’s love’), and 18 items that examine attachment avoidance (‘I don’t feel comfortable opening up to romantic partners’). The participants rate each of the items on a 7-point Likert scale, where 1 = ‘strongly disagree’ and 7 = ‘strongly agree’. Item responses are averaged (after reverse-scoring appropriate items) separately for each subscale to produce a mean score for attachment anxiety and attachment avoidance, with higher scores denoting greater attachment insecurity. This dimensional scoring is in line with research indicating that adult attachment styles are best conceptualised as dimensional constructs (see ^68^). Cronbach’s alphas were α = .93 for anxiety and α = .93 for avoidance in the current sample (across countries).

The TEAQ is a 57-item self-report measure to measure individual experiences, attitudes, and beliefs of physical touch in a variety of contexts (e.g., frequency of touch on a daily basis, cultural and societal influences on attitudes towards touch, previous experience with touch and the potential impact on current attitudes). This questionnaire includes a five-point Likert-type scale (1 = strongly disagree to 5 = strongly agree), with the mean score calculated for each six subscales: Friends and family touch (FFT), Current intimate touch (CIT), Childhood touch (ChT), Attitude to self-care (ASC), Attitude to intimate touch (AIT), and Attitude to unfamiliar touch (AUT). The TEAQ was developed and validated in UK samples and has recently been validated in a SA sample ^61^, indicating its suitability for the current research. Cronbach’s alphas were α = .84 for FFT, α = .88 for CIT, α = .92 for ChT, α = .77 for ASC, α = .89 for AIT, and α = .74 for AUT.

### Statistical Analyses

Our plan of analysis was pre-registered; where we deviated from this initial plan, we state so clearly below. Assumptions of normality were checked using gg-plots, histograms, and tests of normality (e.g., Kolmogorov-Smirnov tests). Continuous covariates were mean centred to avoid multicollinearity issues if they were highly correlated. Individuals with extreme values in group comparisons (i.e. 2.5SD from the group mean in normal or normalised distributions, or else the equivalent Interquartile range) were removed as outliers. No outliers needed to be removed for the touch ratings.

#### Self-report ratings

We planned to carry out a confirmatory factor analysis on touch ratings. If the model fit indicated that they loaded onto one factor, a latent “touch perception” outcome variable would be specified. We also noted that it would be possible that certain ratings (pleasantness, comfort, liking, wanting) but not intensity would load onto one factor. If so, a latent outcome variable would be specified for those ratings that load onto one factor, with other outcomes examined separately. If model fit was poor, we planned to examine ratings separately with corrections for multiple testing applied.

#### Cortical oscillations

Amplitude changes of cortical oscillations were examined by exporting relative power (ERD/S) in theta (4–7 Hz), alpha (8–13 Hz), beta (16–24 Hz) frequency bands from bilateral electrode sites of interest. To restrict the number of comparisons, only electrode clusters associated with touch processing were included: frontal (F1, F2, F3, F4), central (C1, C2, C3, C4, Cz) and parietal (P1, P2, P3, P4, Pz) regions for alpha and beta band power, and frontal and central clusters for theta band power. Frequency bands and electrode clusters were selected based on previous literature showing maximal changes in these bands at the respective sites for tactile stimulation ^29,69,70^ and a previous affective touch study ^32^. Based on an updated review of this literature, some aspects of our approach deviated from our preregistration: frequency bands for analysis deviated from pre-registration in theta (4–7 Hz; 4–8 Hz in preregistration) and beta bands (16–24 Hz; 13–30 Hz in preregistration); relative power was not exported from delta or gamma bands, the former being primarily associated with slow-wave sleep (e.g., reviewed in ^71^), while the latter is frequently produced by microsaccades and muscle activity ^72^; prefrontal, temporal and occipital regions were not included to restrict the number of comparisons; and ERPs are not reported here. Following EEG data cleaning, where <30 trials were accepted for statistical analyses, the condition block was removed from further processing, based on previous work (^27^; see Supplementary Materials).

#### Hypothesis testing

Multivariate linear mixed models (MLMMs) in Stata 18 ^73^ were used to test hypotheses. All data provided by participants was included for touch ratings. MLMMs can handle missing data under the assumption that data is missing at random. Where EEG data was missing entirely (e.g., due to technical issues or persistent noise in the recording), self-report data was still included. Outcomes were self-report ratings and spectral power. In each of these analyses, we included measures of touch experiences and attitudes to touch and adult attachment style as fixed-effect covariates, if they were related to outcome variables. Fixed effects of interest were touch velocity (3cms ^−1^ vs. 18cms ^−1^; within-subjects categorical predictor), and region (arm vs. palm; within-subjects categorical predictor). Including random slopes for velocity and location did not change any of the results, so these were not included in final models. All interaction terms were included, and significant interactions were followed up by specifying Bonferroni-corrected planned contrasts. The intercept of the participant ID was a random effect. We had originally planned to include frequency band and region in analyses as fixed effects of interest. However, we instead ran analyses separately for each frequency band and region, as directly comparing frequency bands or regions was not deemed useful. To account for the increase in number of analyses, we applied Benjamini-Hochberg corrections to the results (false discovery rate set to 5%) and report in the text only those effects that survived correction; full results can be found in the tables.

#### Exploratory analyses

As well as our planned analyses, the above MLMMs were re-run adding country (UK, SA) as a between-subjects categorical predictor and examining its interactions with touch velocity and body region on outcome ratings. Furthermore, we examined the relationships between experiences and attitudes to touch and attachment style (correlation analysis) and demographic variables across countries. Associations between country and demographic variables were investigated, as well as the effect of country (UK, SA) on experiences and attitudes to touch and attachment style, controlling for relevant demographic variables (using ANCOVA or in case of non-normal distributions, bootstrapped regression analyses).

## Data availability

Open Science Framework.

## Acknowledgements

SB was a CIFAR Azrieli Global Scholar in the Brain, Mind and Consciousness Program and is funded by the Oppenheimer Memorial Trust and the National Research Foundation of SA (grant number: 150745). VW received research funding from the Oppenheimer Memorial Trust Award (OMT Ref.2150701). ML is supported by the National Institute for The Humanities and Social Sciences and by a postdoctoral fellowship from the Department of Science and Innovation and National Research Foundation Centre of Excellence in Human Development at the Witwatersrand, Johannesburg, South Africa. CK received research funding from the University of Liverpool.

## Ethics declarations

To conduct this study and collect data, consent was obtained from the Institute of Population Health Ethics Committee, University of Liverpool, and the Human Research Ethics Committee (non-medical) from the University of the Witwatersrand.

