## Supplementary Materials for "Is cultural context the crucial touch? Neurophysiological and self-reported responses to affective touch in women in South Africa and the United Kingdom"

Supplementary Table 1. *Descriptive statistics for touch ratings across countries.*

| | | | Mean | SD | N | Min | Max | Range |
|---|---|---|---|---|---|---|---|---|
| Palm | Slow | Like | 71.81 | 24.45 | 36 | 21 | 100 | 79 |
| | | Want | 60.72 | 28.46 | 36 | 0 | 100 | 100 |
| | | Intense | 35.50 | 30.85 | 36 | 0 | 100 | 100 |
| | | Comfortable | 78.42 | 21.61 | 36 | 2 | 100 | 98 |
| | | Pleasant | 77.81 | 19.56 | 36 | 28 | 100 | 72 |
| | Fast | Like | 67.47 | 25.95 | 36 | 13 | 100 | 87 |
| | | Want | 57.44 | 27.10 | 36 | 5 | 100 | 95 |
| | | Intense | 39.00 | 31.49 | 36 | 0 | 100 | 100 |
| | | Comfortable | 75.78 | 21.76 | 36 | 33 | 100 | 67 |
| | | Pleasant | 70.78 | 22.80 | 36 | 25 | 100 | 75 |
| Arm | Slow | Like | 77.54 | 23.58 | 35 | 16 | 100 | 84 |
| | | Want | 64.57 | 26.03 | 35 | 0 | 100 | 100 |
| | | Intense | 36.74 | 26.94 | 35 | 0 | 85 | 85 |
| | | Comfortable | 79.91 | 24.27 | 35 | 8 | 100 | 92 |
| | | Pleasant | 80.26 | 20.66 | 35 | 14 | 100 | 86 |
| | Fast | Like | 65.67 | 27.00 | 36 | 10 | 100 | 90 |
| | | Want | 57.11 | 25.36 | 36 | 0 | 100 | 100 |
| | | Intense | 37.94 | 30.59 | 36 | 0 | 100 | 100 |
| | | Comfortable | 74.11 | 22.14 | 36 | 30 | 100 | 70 |
| | | Pleasant | 70.31 | 21.08 | 36 | 29 | 100 | 71 |

*Touch ratings:*

As pre-registered, we carried out a confirmatory factor analysis on the ratings, specifying a model with four latent variables (the four conditions), each with five indicators (the five ratings). The specified model did not fit well, $\chi 2(164) = 521.176$, $p < .001$; RMSEA = .25; CFI = .62; SRMR = .164. We next examined Cronbach's alphas for each condition both across countries and within countries. Intensity was negatively correlated with the other ratings (see also Table 3), all alphas improved once intensity was removed, and alphas for the four-item scale were all excellent (see Supplementary Table 2). Theoretically, intensity refers more to discriminative and sensory than affective or hedonic aspects of touch, whereas the latter are captured by evaluative ratings of liking, wanting, and rating how comfortable and pleasant the touch is perceived to be.

Supplementary Table 2. *Cronbach's alpha for all five ratings vs. ratings excluding intensity.*

| | | SA | | UK | | Across countries | |
|---|---|---|---|---|---|---|---|
| | | α | α without intensity | α | α without intensity | α | α without intensity |
| Palm | Slow | 0.75 | 0.88 | 0.80 | 0.93 | 0.75 | 0.90 |
| | Fast | 0.84 | 0.95 | 0.86 | 0.95 | 0.85 | 0.92 |
| Arm | Slow | 0.80 | 0.90 | 0.82 | 0.90 | 0.86 | 0.96 |
| | Fast | 0.75 | 0.89 | 0.88 | 0.96 | 0.80 | 0.94 |

For evaluative touch ratings, across countries, slow touch was evaluated significantly more positively ($M$ = 73.46, $SE$ = 2.70) than fast touch ($M$ = 67.37, $SE$ = 2.70), though fast touch was still rated moderately positively (see Supplementary Table 3 for model results), in line with our hypothesis. There was no significant effect of body region (i.e., whether touch was administered to the arm or the palm), and no significant velocity by body region interaction. There were also no significant findings when intensity was the outcome.

Supplementary Table 3. *Linear mixed modelling results for evaluative (multivariate) and intensity (univariate) touch rating outcomes across countries.*

| | | Evaluation | | | | | Intensity | | | | |
|---|---|---|---|---|---|---|---|---|---|---|---|
| | | *b* | *SE* | *p* | 95% CIs | | *b* | *SE* | *p* | 95% CIs | |
| *Predictors of interest* | Velocity (fast is reference) | 7.90 | 1.98 | **< .001** | 4.01 | 11.79 | -2.09 | 3.12 | .502 | -8.20 | 4.01 |
| | Body region (arm is reference) | 1.07 | 1.97 | .586 | -2.78 | 4.92 | 1.06 | 3.09 | .732 | -4.99 | 7.11 |
| | Velocity × body region | -3.58 | 2.79 | .200 | -9.05 | 1.89 | -1.41 | 4.39 | .748 | -10.00 | 7.19 |
| *Covariates* | Attachment avoidance | -0.49 | 3.49 | .887 | -7.33 | 6.34 | 6.92 | 5.59 | .216 | -4.05 | 17.88 |
| | Attachment anxiety | -7.75 | 3.02 | .010 | -13.68 | -1.83 | -8.94 | 4.85 | .065 | -18.45 | 0.56 |
| | Friends and family touch | 3.96 | 4.65 | .395 | -5.16 | 13.07 | 0.81 | 7.47 | .914 | -13.82 | 15.44 |
| | Current intimate touch | -5.68 | 5.96 | .340 | -17.35 | 5.99 | -11.60 | 9.56 | .225 | -30.34 | 7.13 |
| | Childhood touch | 5.56 | 3.29 | .091 | -0.89 | 12.01 | 6.22 | 5.28 | .239 | -4.13 | 16.57 |
| | Attitude to self-care | -0.48 | 3.29 | .884 | -6.92 | 5.97 | 0.34 | 5.28 | .948 | -10.00 | 10.69 |
| | Attitude to intimate touch | 1.26 | 7.55 | .867 | -13.54 | 16.07 | 11.87 | 12.12 | .328 | -11.89 | 35.63 |
| | Attitude to unfamiliar touch | -1.66 | 3.65 | .649 | -8.81 | 5.49 | 3.24 | 5.85 | .580 | -8.23 | 14.71 |
| *Intercept* | | 78.74 | 26.84 | .003 | 26.13 | 131.35 | 1.33 | 43.07 | .975 | -83.09 | 85.74 |

*Note*. These analyses did not include country as a predictor variable.

*Electroencephalography:*

Following data cleaning, blocks with <30 trials were removed from further processing. No blocks were removed for the UK dataset. Eight blocks were removed from the SA dataset (2 participants with 2 blocks with <30 accepted trials, T19 and T23 in slow palm and fast arm conditions; 4 participants with 1 block with <30 accepted trials, T22, T25, T27 and T34 in fast arm, slow palm, slow arm and slow palm conditions, respectively).

*ERD/S findings across countries*

Grand average time-frequency plots from sensorimotor electrodes showed band power increases focused between 3–6 Hz around the onset of touch, followed by band power decreases concentrated from 4–27 Hz throughout the touch block (Supplementary Figure 1). Visual inspection of topographic plots (Supplementary Figure 2) showed a similar distribution of cortical activation over sensorimotor regions in alpha and beta frequency bands across countries, with the strongest bilateral ERD for touch on the palm, compared to more focal contralateral ERD for touch on the arm.

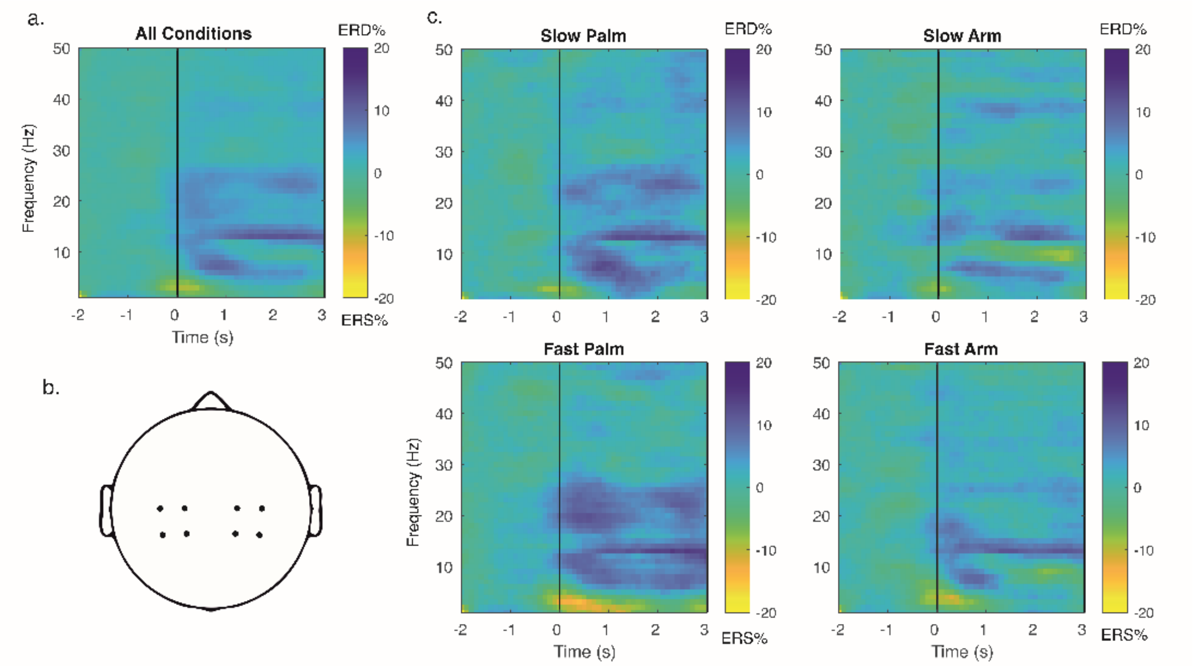


Supplementary Figure 1. *Grand-average time-frequency plots across all conditions (a, c) from electrodes over central and parietal regions of the scalp (b). Data is plotted averaged over all participants*

*with complete datasets (N=28). Colour bar indicates percentage power change from baseline, with positive values indicating ERD, and negative values indicating ERS.*

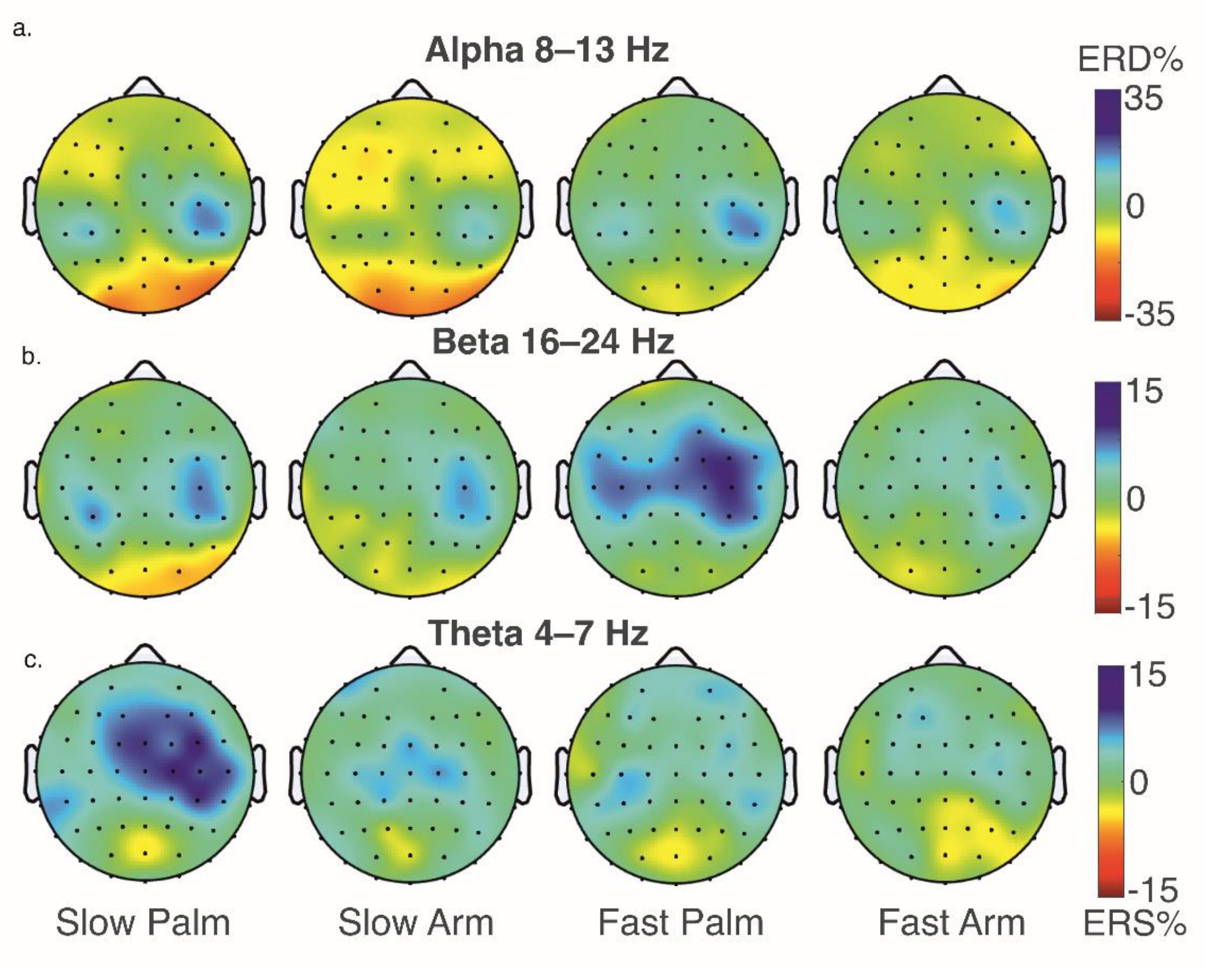


Supplementary Figure 2. *Grand-averaged ERD/S over the entire touch duration (0-3 s) in (a) alpha, (b) beta and (c) theta bands in each of the four conditions, averaged over all participants with complete datasets (N=28). Topographic plots show only electrodes which were common across countries.*

**Findings across countries**

*Alpha*: Alpha ERD at central sites was significantly reduced for slow-velocity (*M* = -.34, *SE* = 3.48) compared to faster-velocity (*M* = 5.42, *SE* = 3.47) touch across body regions (i.e., arm or palm; see Supplementary Figures 3a, c). Moreover, alpha ERS was significantly greater for the arm (*M* = -.70, *SE* = 3.47) vs. palm (*M* = 5.87, *SE* = 3.47) at central and parietal sites (*M* = -5.48, *SE* = 1.76 for palm; *M* = -9.00, *SE* = 1.76 for arm) across stroking speeds (Supplementary Figure 3d, e; Supplementary Table 4).

These findings are very similar to those reported in the main text, but please note this model did not include country as a predictor variable.

*Beta*: As for alpha, central beta ERD was significantly greater for the palm (*M* = 6.19, *SE* = 1.46) compared to the arm (*M* = 1.76, *SE* = 1.46 for arm; Supplementary Figure 3b, f), as expected. However, this body region effect was further qualified by a significant velocity by body region interaction (Supplementary Table 4). Planned contrasts showed stronger ERD for fast (*M* = 8.53, *SE* = 1.54) vs. slow touch (*M* = 3.77, *SE* = 1.56) at the palm (contrast = 4.75, *SE* = 1.04, $p < .001$) but not the arm region (fast arm: *M* = 1.72, *SE* = 1.56; slow arm: *M* = 1.80, *SE* = 1.54; contrast = -.07, *SE* = .104, *p* = .944), in line with our hypotheses.

*Theta*: Contrary to beta body region effects which were evident for the faster touch to the palm, the interaction between velocity and body region for frontal theta diverged for arm and palm: At the arm, ERD was greater for fast (*M* = 3.92, *SE* = 2.43) than slow (*M* = 1.20, *SE* = 2.41) touch (contrast = 2.72, *SE* = 1.31, *p* = .037), in contrast to von Mohr et al. (2018), while for the palm, ERD was reduced for fast (*M* = 4.88, *SE* = 2.40) vs. slow (*M* = 7.63, *SE* = 2.43) touch (contrast = -2.75, *SE* = 1.31, *p* = .035), in line with Kraus et al. (2020); see Supplementary Table 4 and Supplementary Figure 2c.

Supplementary Table 4. *Multivariate linear mixed modelling results for alpha, beta, and theta ERD/S across countries.*

| | | | *Central electrode sites* | | | | | *Parietal electrode sites* | | | | |
|---|---|---|---|---|---|---|---|---|---|---|---|---|
| **Frequency band** | | **Predictor** | **b** | **SE** | **p** | **95% CIs** | | **b** | **SE** | **p** | **95% CIs** | |
| **Alpha** | *Predictors of interest* | Velocity (fast is reference) | -7.11 | 1.96 | **<.001** | -10.96 | -3.26 | -0.82 | 1.67 | **.001** | -4.09 | 2.44 |
| | | Body region (arm is reference) | 5.24 | 1.94 | **.007** | 1.43 | 9.05 | 5.90 | 1.65 | **.003** | 2.67 | 9.14 |
| | | Velocity × body region | 2.69 | 2.80 | .337 | -2.80 | 8.17 | -4.85 | 2.37 | .869 | -9.50 | -0.20 |
| | *Covariates* | Attachment avoidance | 8.68 | 4.85 | .073 | -0.82 | 18.18 | 7.55 | 2.36 | .712 | 2.92 | 12.18 |
| | | Attachment anxiety | -10.63 | 4.41 | .016 | -19.27 | -1.99 | -6.28 | 2.14 | .454 | -10.48 | -2.07 |
| | | Friends and family touch | 0.58 | 6.02 | .923 | -11.22 | 12.38 | -0.48 | 2.93 | .773 | -6.22 | 5.25 |
| | | Current intimate touch | -10.60 | 7.60 | .163 | -25.50 | 4.30 | -1.37 | 3.72 | .535 | -8.66 | 5.92 |
| | | Childhood touch | -3.79 | 4.19 | .365 | -12.00 | 4.42 | 1.53 | 2.04 | .086 | -2.47 | 5.52 |
| | | Attitude to self-care | -6.59 | 4.36 | .130 | -15.13 | 1.95 | 0.61 | 2.12 | .029 | -3.54 | 4.77 |
| | | Attitude to intimate touch | 4.08 | 10.15 | .688 | -15.81 | 23.97 | 3.08 | 4.96 | **.001** | -6.64 | 12.80 |
| | | Attitude to unfamiliar touch | 10.76 | 4.90 | .028 | 1.16 | 20.35 | 4.09 | 2.38 | **.003** | -0.58 | 8.76 |

| | | | | | | | | | | | | |
|---|---|---|---|---|---|---|---|---|---|---|---|---|
| | *Intercept* | | 35.10 | 34.62 | .311 | -32.76 | 102.95 | -37.15 | 16.98 | .869 | -70.43 | -3.88 |
| **Beta** | *Predictors of interest* | Velocity (fast is reference) | 0.07 | 1.04 | .944 | -1.96 | 2.11 | -0.82 | 1.01 | .416 | -2.79 | 1.16 |
| | | Body region (arm is reference) | 6.80 | 1.03 | **<.001** | 4.79 | 8.81 | 2.23 | 1.00 | .025 | 0.28 | 4.19 |
| | | Velocity × body region | -4.82 | 1.48 | **.001** | -7.72 | -1.93 | -1.51 | 1.44 | .294 | -4.32 | 1.31 |
| | *Covariates* | Attachment avoidance | 1.45 | 2.01 | .470 | -2.49 | 5.40 | 3.36 | 2.04 | .099 | -0.64 | 7.36 |
| | | Attachment anxiety | -1.68 | 1.83 | .359 | -5.27 | 1.91 | -3.76 | 1.85 | .043 | -7.40 | -0.13 |
| | | Friends and family touch | -2.71 | 2.50 | .279 | -7.61 | 2.19 | -2.58 | 2.53 | .309 | -7.54 | 2.39 |
| | | Current intimate touch | -1.05 | 3.16 | .740 | -7.25 | 5.15 | 0.01 | 3.20 | .997 | -6.27 | 6.29 |
| | | Childhood touch | -0.40 | 1.74 | .817 | -3.81 | 3.01 | -1.42 | 1.76 | .422 | -4.87 | 2.04 |
| | | Attitude to self-care | 0.42 | 1.81 | .815 | -3.12 | 3.97 | 1.61 | 1.83 | .380 | -1.98 | 5.20 |
| | | Attitude to intimate touch | -0.32 | 4.22 | .94 | -8.59 | 7.95 | -0.54 | 4.27 | .899 | -8.92 | 7.83 |
| | | Attitude to unfamiliar touch | 5.91 | 2.03 | **.004** | 1.92 | 9.89 | 6.48 | 2.06 | **.002** | 2.44 | 10.52 |
| | *Intercept* | | -0.01 | 14.41 | .999 | -28.25 | 28.24 | -7.73 | 14.59 | .597 | -36.33 | 20.88 |
| **Theta** | | ***Frontal electrode sites*** | | | | | | ***Central electrode sites*** | | | | |

| | | | | | | | | | | | |
|---|---|---|---|---|---|---|---|---|---|---|---|
| *Predictors of interest* | Velocity (fast is reference) | -2.70 | 1.31 | .039 | -5.26 | -0.14 | 0.82 | 1.46 | .573 | -2.03 | 3.67 |
| | Body region (arm is reference) | 0.98 | 1.29 | .449 | -1.56 | 3.51 | 0.55 | 1.44 | .703 | -2.28 | 3.37 |
| | Velocity × body region | 5.40 | 1.86 | **.004** | 1.75 | 9.05 | 2.19 | 2.07 | .291 | -1.87 | 6.25 |
| *Covariates* | Attachment avoidance | 4.04 | 2.49 | .104 | -0.83 | 8.92 | 4.27 | 2.36 | .070 | -0.35 | 8.90 |
| | Attachment anxiety | -4.37 | 2.26 | .053 | -8.80 | 0.06 | -2.90 | 2.14 | .176 | -7.10 | 1.30 |
| | Friends and family touch | 3.57 | 3.09 | .248 | -2.48 | 9.62 | 4.16 | 2.92 | .155 | -1.58 | 9.89 |
| | Current intimate touch | -6.30 | 3.90 | .107 | -13.95 | 1.35 | -6.93 | 3.71 | .062 | -14.20 | 0.34 |
| | Childhood touch | -1.10 | 2.15 | .608 | -5.31 | 3.11 | 0.62 | 2.04 | .762 | -3.38 | 4.61 |
| | Attitude to self-care | -4.13 | 2.23 | .064 | -8.51 | 0.25 | -6.35 | 2.12 | **.003** | -10.50 | -2.20 |
| | Attitude to intimate touch | 0.53 | 5.21 | .920 | -9.69 | 10.74 | 1.33 | 4.95 | .788 | -8.36 | 11.03 |
| | Attitude to unfamiliar touch | 6.13 | 2.51 | .015 | 1.21 | 11.05 | 5.55 | 2.38 | .020 | 0.89 | 10.21 |
| *Intercept* | Constant | 13.23 | 17.80 | .457 | -21.65 | 48.11 | 8.41 | 16.91 | .619 | -24.74 | 41.56 |

*Note*. *p* values that survived the Benjamini-Hochberg correction are highlighted in bold font. Country was not included as a predictor in these models.

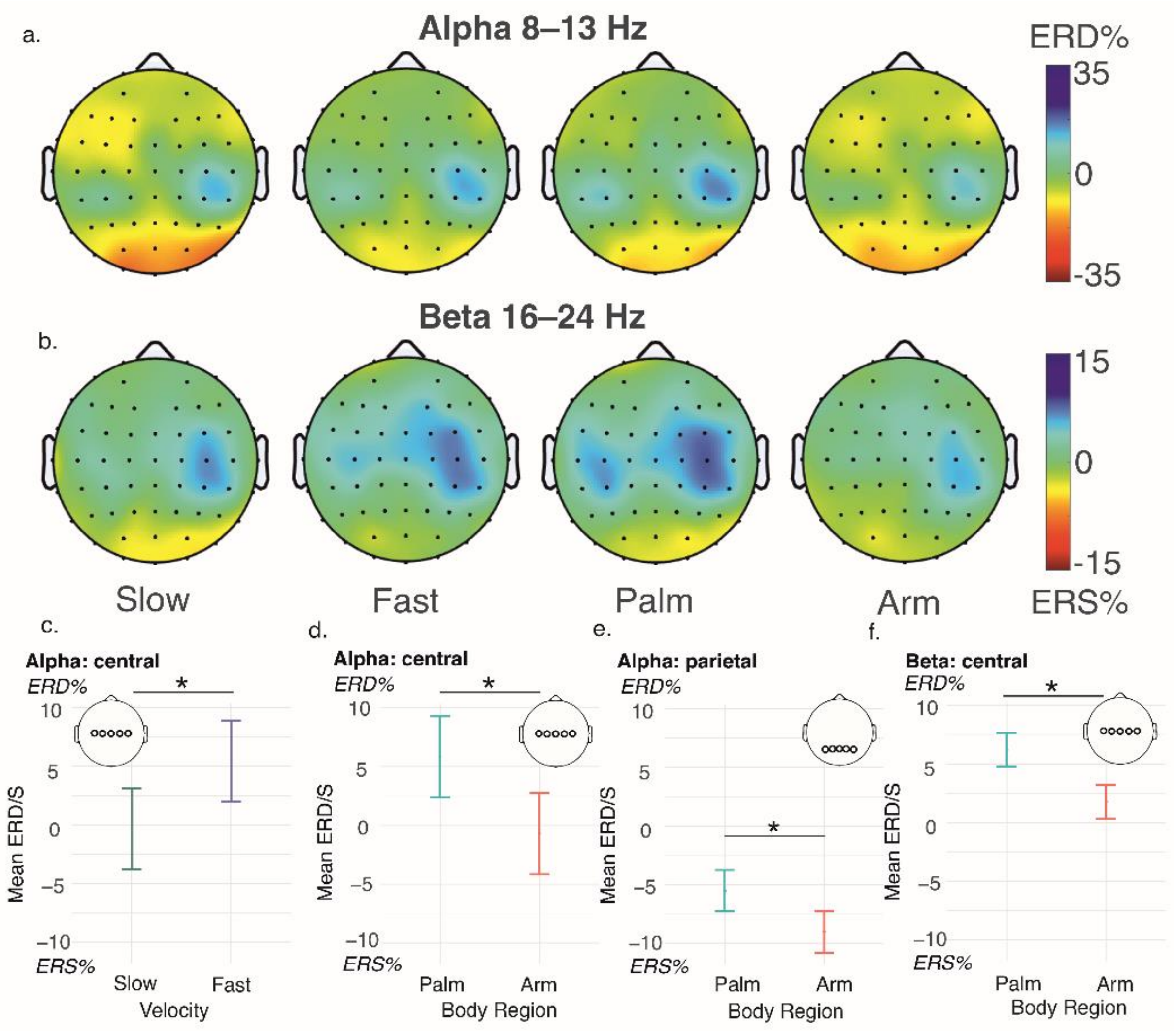


Supplementary Figure 3. *Effects of touch velocity and body region on alpha (a) and beta (b) event-related synchronisation/desynchronisation (ERS/ERD) over the entire touch duration (0-3 s). Mean ERD/S for planned comparisons are plotted over central sites in alpha (c, d) and beta (f) bands, and in parietal sites in the alpha band (e). Error bars denote ±1 standard error of the mean.*
